# Small RNAs activate *Salmonella* pathogenicity island 1 by modulating mRNA stability through the *hilD* mRNA 3′ UTR

**DOI:** 10.1101/2022.09.07.507058

**Authors:** Sabrina Z. Abdulla, Kyungsub Kim, Muhammad S. Azam, Yekaterina A. Golubeva, Fatih Cakar, James M. Slauch, Carin K. Vanderpool

## Abstract

*Salmonella enterica* serovar Typhimurium is an enteric pathogen associated with food-borne disease. *Salmonella* invades the intestinal epithelium using a type three secretion system encoded in *Salmonella* pathogenicity Island 1 (SPI-1). SPI-1 genes are tightly regulated by a complex feed-forward loop to ensure proper spatial and temporal expression. Most regulatory input is integrated at HilD, through control of *hilD* mRNA translation or HilD protein activity. The *hilD* mRNA possesses a 310-nucleotide 3′ untranslated region (UTR) that influences HilD and SPI-1 expression, and this regulation is dependent on Hfq and RNase E, cofactors known to mediate small RNA (sRNA) activities. Thus, we hypothesized that the *hilD* mRNA 3′ UTR is a target for sRNAs. Here we show that the sRNAs, SdsR and Spot 42 regulate SPI-1 by targeting different regions of the *hilD* mRNA 3′ UTR. Regulatory activities of these sRNAs depend on Hfq and RNase E, in agreement with previous roles found for both at the *hilD* 3′ UTR. We show that SdsR and RNase E are responsible for the accumulation of variable fragments of the *hilD* mRNA 3′ UTR. Collectively, this work suggests that these sRNAs targeting the *hilD* mRNA 3′ UTR regulate *hilD* mRNA levels by interfering with RNase E-dependent mRNA degradation. Our work provides novel insights into mechanisms of sRNA regulation at bacterial mRNA 3′ UTRs and adds to our knowledge of post-transcriptional regulation of the SPI-1 complex feed-forward loop.

**Importance:** *Salmonella* are prominent food-borne pathogens, infecting millions of people a year. To express virulence genes at the correct time and place in the host, *Salmonella* uses a complex regulatory network that senses environmental conditions. Known for their role in allowing quick responses to stress and virulence conditions, we investigate the role of small RNAs in facilitating precise expression of these genes. We provide evidence that the 3′ untranslated region of the *hilD* mRNA, encoding a key virulence regulator, is a target for small RNAs and the ribonuclease RNase E. The small RNAs play a role in stabilizing *hilD* mRNA to allow proper expression of *Salmonella* virulence genes in the host.

## Introduction

The gastrointestinal pathogen, *Salmonella enterica* serovar Typhimurium, invades the host intestinal epithelium to induce a typically self-limiting gastroenteritis (1, 2), but can also cause serious systemic disease (3). *Salmonella* breaches the host intestinal epithelial barrier using a type 3 secretion system (T3SS) that injects effector proteins directly into host cells, resulting in actin rearrangement and induction of an inflammatory response (4–6). The T3SS structural genes, along with several effectors needed for invasion, are encoded on *Salmonella* pathogenicity island 1 (SPI-1). Loss of SPI-1 prevents *Salmonella* from invading host cells during infection, resulting in strong virulence defects. Conversely, SPI-1 overexpression negatively affects *Salmonella* fitness (5, 7). To ensure precise control, SPI-1 gene expression is regulated by a complex network that responds to many signals (8, 9).

HilA activates transcription of the SPI1 structural genes. Transcription of *hilA* is regulated by three AraC-like proteins, HilD, HilC, and RtsA. These factors form a complex feed-forward loop in which they regulate their own and each other’s expression along with expression of *hilA* (8, 9) (Fig. 1). Environmental signals regulate SPI-1 by integrating into the SPI-1 feed-forward loop at multiple levels (9, 10). Many signals regulate SPI-1 at the level of *hilD* translation or HilD protein activity (9, 11, 12). There are only a few examples of signal integration at *hilC* or *rtsA* (27). Instead, HilC and RtsA act primarily to amplify signals that control HilD levels (9, 13, 14). In recent decades, small RNAs (sRNAs) have emerged as important post-transcriptional regulators that commonly regulate virulence and stress response regulatory circuits (15, 16). Through base-pairing interactions with mRNA targets, sRNAs can regulate translation or mRNA stability quickly and efficiently. The RNA chaperones Hfq and ProQ aid with sRNA stability and/or facilitate target binding (17–20). sRNAs can either positively or negatively regulate targets (21, 22). The RNase E degradosome often plays an important role in sRNA-mediated regulation and a common mechanism of positive regulation by sRNAs involves blocking RNase E-mediated degradation of target transcripts (23). Conversely, sRNAs can repress translation and directly or indirectly recruit RNase E to promote target degradation (24). There is clear evidence for sRNA-mediated post-transcriptional regulation of *hilD* by mechanisms involving both its 5′ and 3′ untranslated regions (UTRs) (8, 10, 16, 25–31).

**Figure 1.**
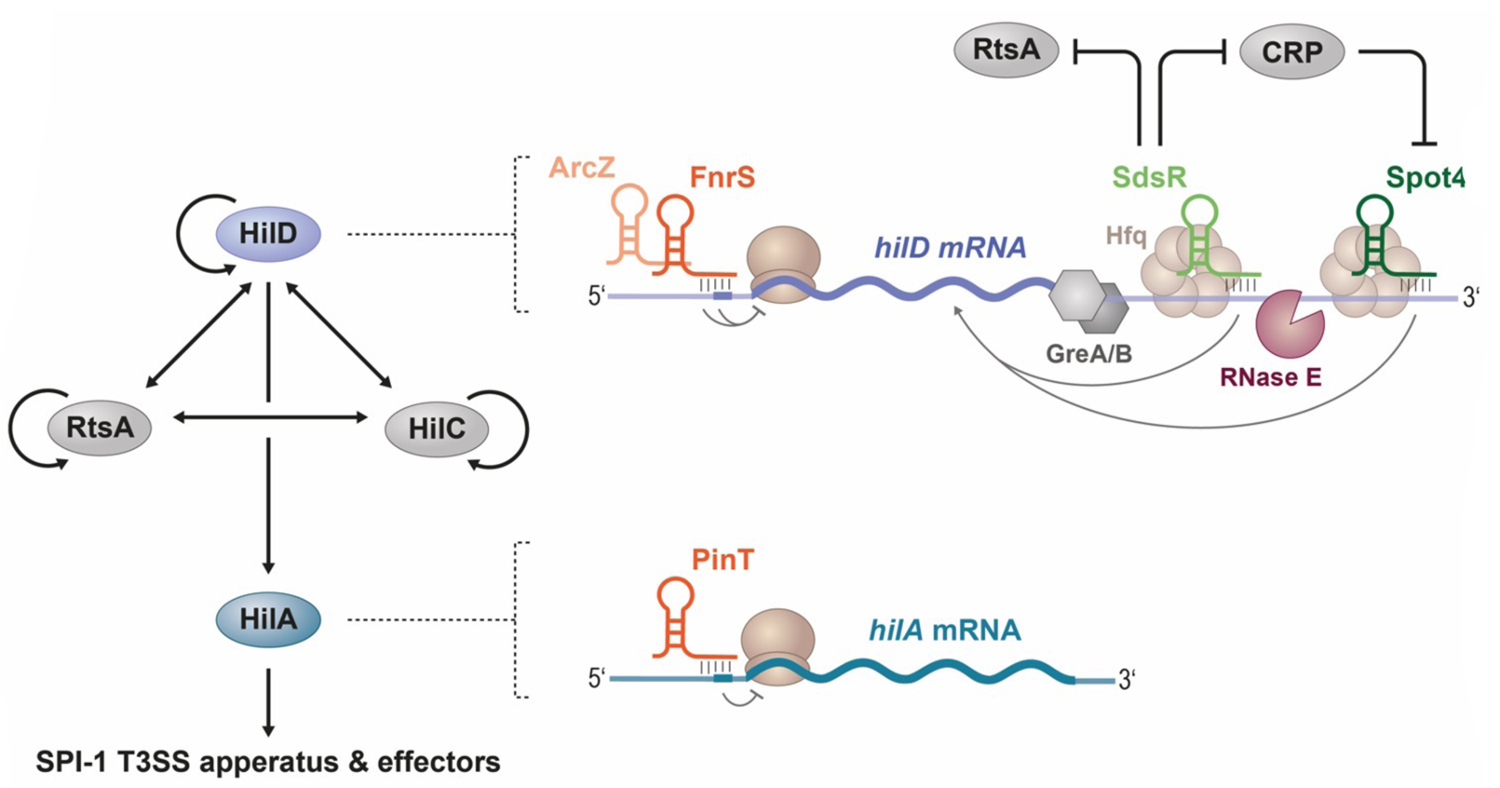
*Salmonella* pathogenicity island 1 (SPI-1) is regulated by a complex feed-forward loop. *Salmonella* pathogenicity island I (SPI-1) genes are regulated by a complex feed-forward loop consisting of three AraC-like regulators, HilD, HilC, and RtsA. These three regulators contribute to production of a fourth AraC-like transcription factor, HilA, which activates expression of SPI-1 type 3 secretion system apparatus and effector genes. Regulation of SPI-1 at a post-transcriptional level by small RNAs (sRNAs) also contributes to control of *Salmonella* invasion and pathogenesis. Both *hilD* and *hilA* mRNAs are targets of sRNA-mediated regulation. The *hilD* mRNA 3′ UTR is 310-nt long and is targeted by RNase E. We propose that during SPI-1 inducing conditions, sRNA binding at the *hilD* mRNA 3′ UTR protects the *hilD* mRNA 3’UTR from RNase E, thus stabilizing *hilD* mRNA. Previous work demonstrated that *hilD* and *hilA* mRNA 5’ UTRs are also targets for sRNA regulatory inputs.

We have shown that the sRNAs FnrS and ArcZ regulate *hilD* in response to oxygen levels by base pairing with sequences in the *hilD* 5′ UTR, confirming a role for sRNAs in integration of environmental signals into the SPI-1 regulatory network (28). Additionally, the sRNA PinT was shown to regulate both *hilD* and *rtsA* (27), suggesting that sRNAs mediate control of SPI-1 at multiple levels. While sRNAs often regulate translation and mRNA stability through interactions with mRNA 5′ UTRs and coding regions of genes, López-Garrido, *et al.,* determined that the 310-nucleotide (nt) long *hilD* 3′ UTR is a regulatory element that affects levels of *hilD* mRNA (25). Deletion of the *hilD* mRNA 3′ UTR dramatically increases levels of *hilD* mRNA, suggesting that it destabilizes the *hilD* mRNA under conditions where SPI-1 is not induced. A similar, but less dramatic increase in *hilD* mRNA levels is seen in RNase E mutants (*rne131*), while levels of *hilD* mRNA are decreased in Δ*hfq* mutants, implicating sRNAs in controlling *hilD* mRNA levels through interactions with the *hilD* 3′ UTR (25). Canonical mechanisms of sRNA regulation often involve blocking or exposing ribosome binding sites; sRNAs base pairing with the *hilD* mRNA 3′ UTR would potentially reveal novel mechanisms of regulation. Recently, El Mouali, *et al.,* found that the sRNA Spot 42 activates SPI-1 by a mechanism involving the *hilD* mRNA 3′ UTR (26), suggesting that there is indeed a role for sRNA regulators at the *hilD* mRNA 3′ UTR.

In this paper, we focus on sRNA regulators that target the *hilD* mRNA 3′ UTR. Using genetic and biochemical approaches, we show that the sRNA SdsR activates *hilD* through direct interactions with the *hilD* 3′ UTR. We find that SdsR and Spot 42 target distinct regions of the *hilD* 3′ UTR, and both sRNAs activate *hilD* via a mechanism involving RNase E. SdsR-dependent changes in RNA structure suggest that SdsR binding to the *hilD* mRNA 3′ UTR alters the accessibility of RNase E cleavage sites, thus stabilizing *hilD* mRNA and promoting SPI-1 gene expression. Further, the effect of these sRNAs is reflected during infection. This work provides new evidence of 3′ UTR-mediated post-transcriptional regulation in bacteria, insight into mechanisms of sRNAs that target 3′ UTRs and broadens the target regulon of SdsR.

## Results

### *The hilD* mRNA 3′ UTR affects HilD expression levels

The *hilD* mRNA contains a 310-nt 3′ UTR that affects *hilD* mRNA levels and is influenced by regulators such as Hfq, RNase E, and Gre factors (20, 25, 32) (Fig. 1). To define regulatory regions within the *hilD* mRNA 3′ UTR, we constructed a series of *lacZ* transcriptional reporter fusions containing different regions of the 3′ UTR. The Δ3′ UTR reporter has *lacZ* fused immediately downstream of the *hilD* stop codon (Fig. 2A). The +90 3′ UTR and +180 3′ UTR have *lacZ* inserted 90 or 180 nts, respectively, downstream of the *hilD* stop codon, while the Full 3′ UTR fusion has *lacZ* inserted 280 nts downstream of the stop codon, just prior to the intrinsic terminator stem-loop (Fig. 2A). Activity of these reporters was tested under high-salt, low-oxygen, SPI-1-inducing conditions (33). The Full 3′ UTR fusion yielded a very low level of activity compared to each of the truncated fusions, while expression from the Δ3′ UTR fusion was increased ∼60-fold (Fig. 2B). The +90 and +180 fusions had intermediate levels of activity, suggesting that *hilD* mRNA levels increase as the 3′ UTR is progressively truncated from the 3′ end (Fig. 2B). These observations support the idea that the 3′ UTR of *hilD* mRNA confers instability and may be targeted by factors such as RNase E, consistent with previous studies (25).

**Figure 2:**
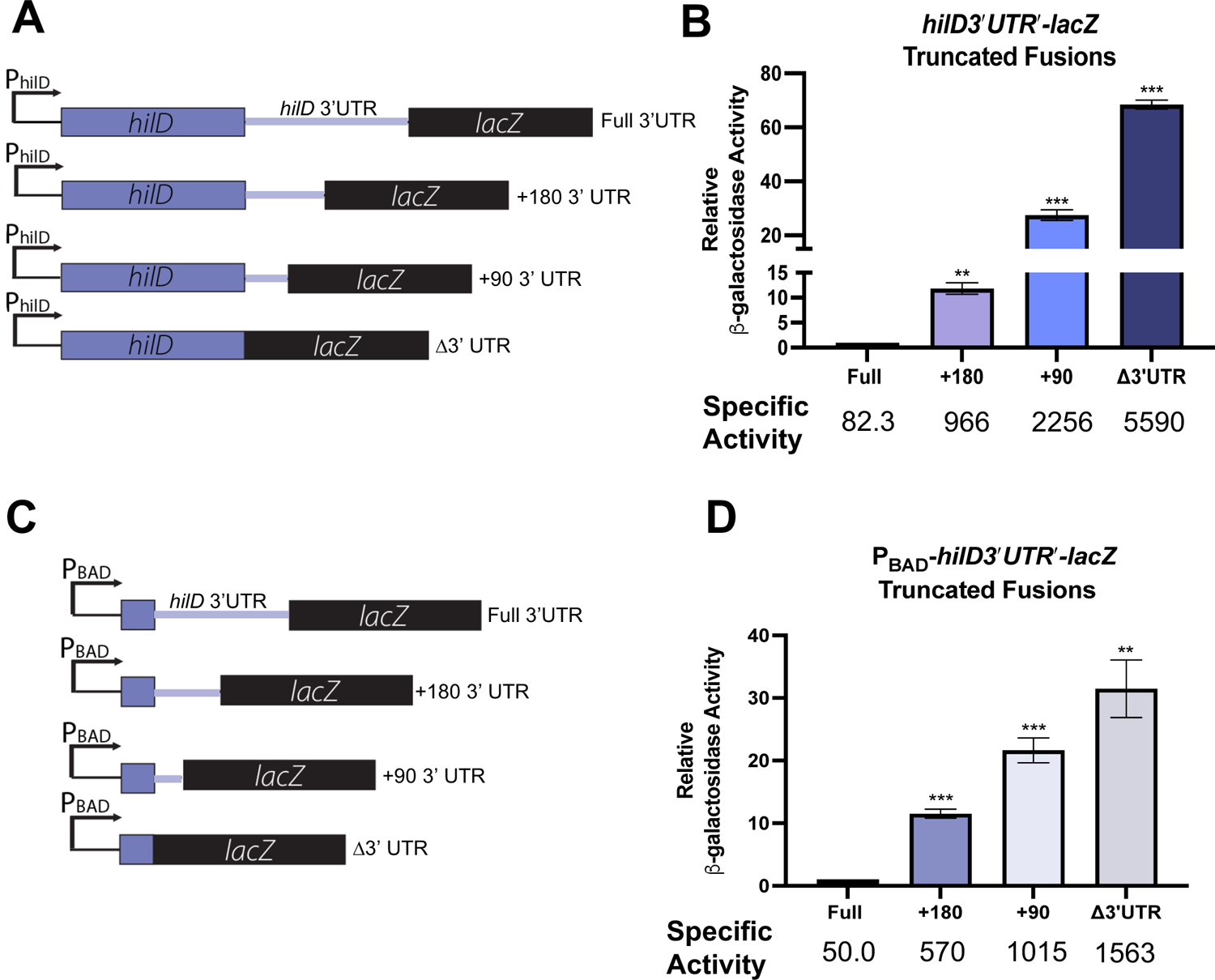
*hilD* mRNA 3’ UTR acts as a regulatory element to regulate levels of *hilD* mRNA. (A) A diagram of *hilD 3*′*UTR*′*-lacZ* fusions in *Salmonella*. *lacZ* is inserted in the *hilD* locus before the *hilD* terminator (Full), after +180 nt of *hilD* mRNA 3’UTR (+180), +90 nt of *hilD* mRNA 3’UTR (+90), or directly after the *hilD* CDS (Δ3′ UTR). (B) *hilD 3*′*UTR*′*-lacZ* fusion strains were grown in no salt LB (NSLB) overnight, then subcultured into high salt LB (HSLB) and grown statically for 18 hours at 37 degrees C to promote SPI-1 activation. Samples were harvested and β-galactosidase assays were performed. (C) Diagram of P_BAD_-*hilD* 3′*UTR’-lacZ* fusions in *E. coli*. Fusions are similar to *Salmonella* fusions in (A), but *hilD 3*′*UTR*′*-lacZ* is inserted downstream of P_BAD_ promoter. See text for a more detailed description of fusion construction. (D) *E. coli hilD 3*′*UTR*′*-lacZ* strains were grown in LB with 0.002% L-arabinose until OD_600_=0.05, then harvested for β-galactosidase assays. Relative β-galactosidase units were obtained by normalizing β-galactosidase activity to WT control and reported as mean ± standard deviation. Error bars are standard deviations of the results of three independent experiments, obtained by unpaired *t* test, n=3. **, *P* < 0.005; ***, *P* < 0.0005.

We next wanted to determine whether the *hilD* mRNA 3′ UTR is necessary and sufficient for the regulatory effects we observed in *Salmonella*. Previous work showed that fusion of the *hilD* mRNA 3′ UTR downstream of *lacZ* or GFP reporters in both *Salmonella* and *E. coli* was sufficient to confer negative regulation of the reporter (25, 34). To further evaluate whether the 3′ UTR-dependent regulatory effects observed for *Salmonella hilD* are dependent on the context at the native locus or *Salmonella*-specific factors such as SPI-1 regulators, we constructed analogous reporter fusions in *E. coli*. These transcriptional fusions contain the region from +660 nt of the *hilD* coding sequence (300 bp upstream of the stop codon) through various lengths of the *hilD* 3′ UTR corresponding to the end points of the fusions in *Salmonella* (Δ3′ UTR: +660-+900, +90: +660-+990, +180: +660-1080, Full: +660-+1237). These regions were inserted into the *E. coli* chromosome under the control of an arabinose-inducible P_BAD_ promoter and 5’ to the *lacZ* gene (32, 35) (Fig. 2C). The *E. coli* fusions showed a similar pattern of increased *hilD* expression with shorter *hilD* mRNA 3′ UTR lengths (Fig. 2D). These data further support results from other groups and our results in *Salmonella* (Figs 2A, B), demonstrating that the *hilD* mRNA 3′ UTR acts as an independent regulatory element. Our data are consistent with the model that the 3′ UTR acts as a destabilizing element that reduces *hilD* mRNA levels.

### RNase E and Hfq modulate HilD expression through the 3′ UTR

We have previously shown that several regulatory signals are integrated into the SPI-1 regulatory network through sRNAs that act at the *hilD* mRNA 5′ UTR and *hilA* mRNA 5′ UTR (9, 27, 28) (Fig. 1). Additionally, recent work has shown that the sRNA Spot 42 acts through the *hilD* mRNA 3′ UTR to regulate SPI-1 (26). Based on previous work showing that Hfq and RNase E play a role in regulation at the *hilD* mRNA 3′ UTR (25), we hypothesized that sRNAs could stabilize *hilD* mRNA by acting with Hfq to occlude RNase E processing sites within the *hilD* mRNA 3′ UTR. To confirm that Hfq and RNase E play a role in regulation of *hilD* via a mechanism involving the 3′ UTR, we measured activity of the Full 3′ UTR fusion in a *Salmonella* !ι*hfq* strain, which lacks the RNA chaperone involved in mediating sRNA-dependent regulation. We also measured fusion activity in a *Salmonella rne131* strain (28), which produces a truncated RNase E protein that cannot assemble the RNA degradosome complex involved in many sRNA-mediated regulatory mechanisms (36). Compared to the *hfq^+^ rne*^+^ (WT) background, the activity of the fusion in the *Δhfq* strain was reduced by ∼50% (Fig. 3). In contrast, fusion activity increased about 3-fold in the *rne131* mutant compared to the WT parent (Fig. 3). Together, these results are consistent with previous work (20, 25) and support our hypothesis that Hfq-dependent sRNAs may protect the *hilD* mRNA 3′ UTR from RNase E-dependent degradation.

**Figure 3:**
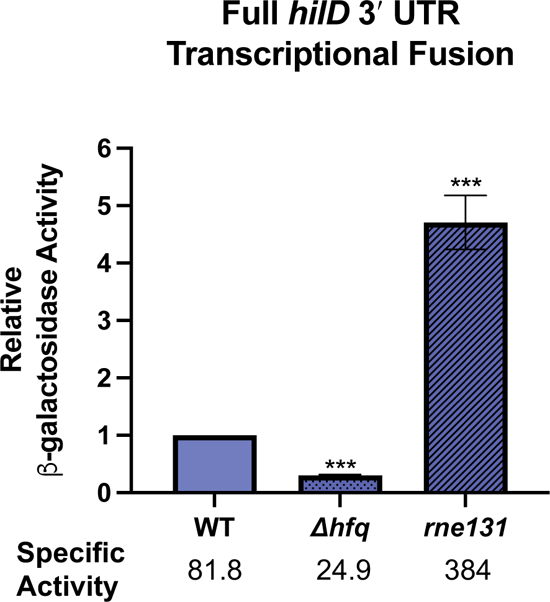
Hfq and RNase E act at the *hilD* mRNA 3′ UTR. *Salmonella strains containing hilD 3*′*UTR*′*-lacZ* transcriptional fusion at the *hilD* locus in wild-type (WT), *rne131* or Δ*hfq* backgrounds. Strains were grown in no salt LB, then subcultured 1:100 into high salt LB. Subcultures were grown statically for 20 hours at 37°C. Relative β-galactosidase units were obtained by normalizing β-galactosidase activity to WT control and reported as mean ± standard deviation. Error bars are standard deviations of the results of three independent experiments, obtained by unpaired *t* test, n=3. ***, *P* < 0.0005.

### sRNAs regulate SPI-1 in a *hilD* mRNA 3′ UTR dependent manner

Spot 42 was previously found to regulate SPI1 via the *hilD* mRNA 3′ UTR (27, 28, 37). We postulated that other sRNA regulators of the *hilD* mRNA 3′ UTR are likely. Using IntaRNA 2.0 to obtain base-pairing predictions between sRNAs and the *hilD* mRNA 3′ UTR and testing a series of *lacZ* gene fusions to confirm regulation at the appropriate levels of SPI-1, we identified SdsR as a candidate regulator acting at the *hilD* 3′ UTR. SdsR and Spot 42 both activated the *hilA′-lacZ* transcriptional fusion by 5- to 6-fold, respectively (Fig. 4A). Neither sRNA substantially regulated a *hilD′-′lacZ* translational fusion (Fig. 4B), or a *hilA′-′lacZ* translational fusion in *E. coli* (Fig. 4C) suggesting that they were not regulating *hilA* expression through interactions at the 5′ end of the *hilD* mRNA or directly at the *hilA* mRNA. In contrast, both SdsR and Spot 42 activated the Full 3′ UTR fusion by 1.5- to 2-fold (Fig. 4D), suggesting that they regulate *hilA* indirectly through activation of *hilD* via the 3′ UTR. Note that the sRNAs increased *hilA* transcriptional fusion activity to a greater degree than the *hilD* reporter, consistent with signal amplification via the complex feed-forward regulatory loop (14). We next expressed SdsR and Spot 42 in the Δ3′ UTR fusion. In contrast with the Full fusion, neither sRNA activated the Δ3′ UTR fusion (Fig. 4D). Instead, both SdsR and Spot 42 slightly repressed the Δ3′ UTR fusion (Fig. 4D). These data demonstrate that both SdsR and Spot 42 regulate SPI-1 primarily by a mechanism involving the *hilD* mRNA 3′ UTR.

**Figure 4:**
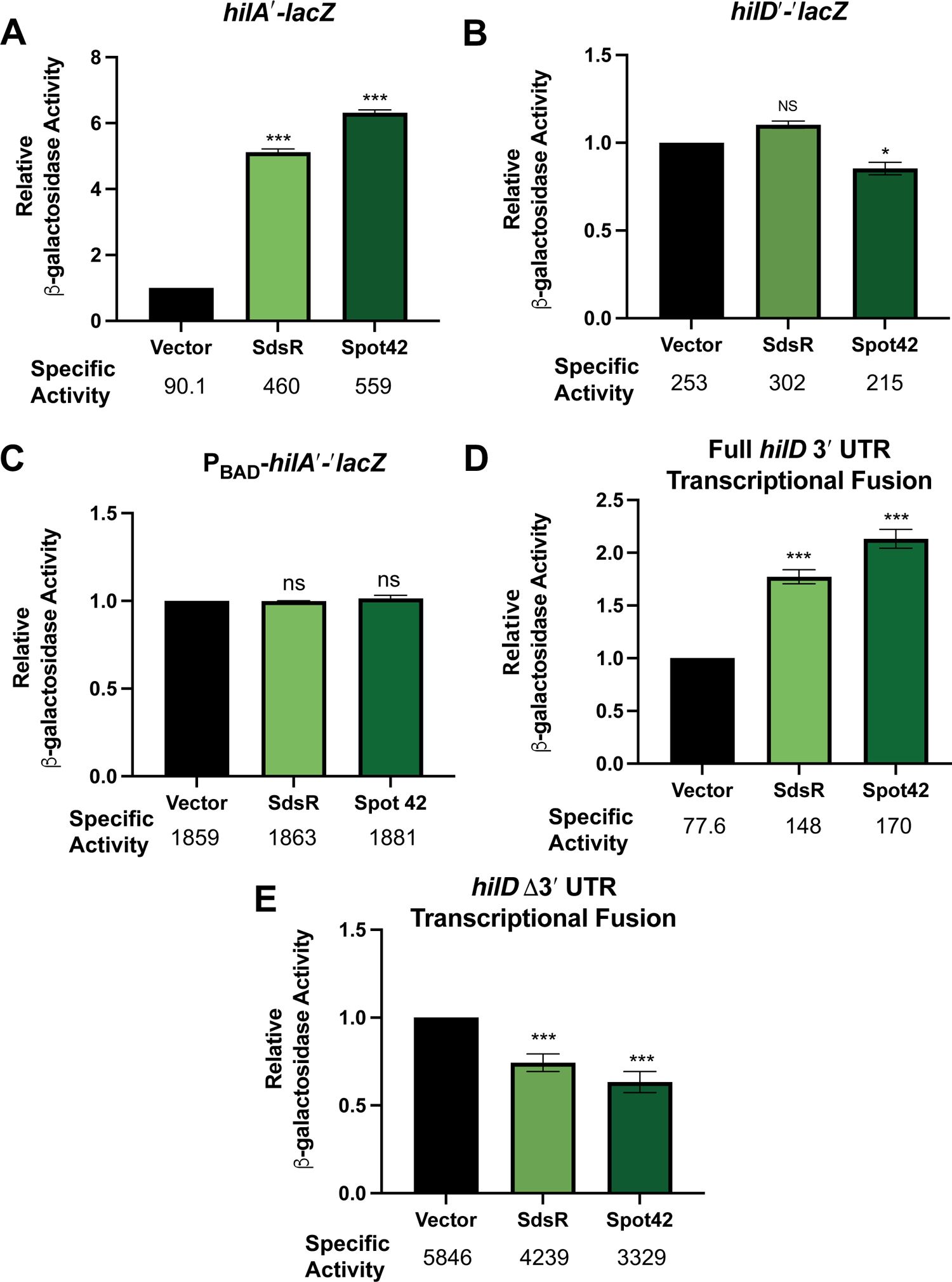
sRNAs regulate SPI-1 in a *hilD* mRNA 3’ UTR-dependent manner. Plasmids encoding SdsR and Spot 42 in *Salmonella* strains containing (A) *hilA*′*-lacZ* transcriptional fusion, (B) *hilD*′*-*′*lacZ* translational fusion, (D) Full *hilD 3*′*UTR*′*-lacZ* transcriptional fusion, and (E) *hilD Δ3*′*UTR*′*-lacZ* transcriptional fusion. Strains were grown in no salt LB, then subcultured 1:100 to high salt LB. Subcultures were grown statically for 20 hours at 37°C. (C) *E. coli* P_BAD_-*hilA*′*-*′*lacZ* strains producing SdsR and Spot 42 were grown in LB with 0.002% L-arabinose until OD_600_=0.05, then harvested for β-galactosidase assays. Relative β-galactosidase units were obtained by normalizing β-galactosidase activity to WT control and reported as mean ± standard deviation. Error bars are standard deviations of the results of three independent experiments, obtained by unpaired *t* test, n=3. *, *P* < 0.05; **, *P* < 0.005; ***, *P* < 0.0005, ns, not significant.

### SdsR regulates SPI-1 at multiple levels

The SPI-1 complex feed-forward loop allows for regulatory inputs at multiple levels. We recently found that the sRNA PinT regulates SPI-1 by at least two mechanisms, regulating translation of both *hilA* and *rtsA* (27). We wondered if the slight repression of the Δ3′ UTR fusion observed for SdsR and Spot 42 was due to negative regulation by these sRNAs at other levels of SPI-1. SdsR has a broad target spectrum and was previously shown to negatively regulate *rtsA* (38). We first used *hilC* and *rtsA* translational fusions (in an *E. coli* host) to determine whether SdsR or Spot 42 directly regulate at other levels of SPI-1. Neither SdsR nor Spot 42 regulated a *hilC′-′lacZ* fusion (Fig. S1A). As shown previously, SdsR and PinT both repress *rtsA′-′lacZ* activity, while Spot 42 had no significant impact on *rtsA* translation (Fig. S1B). Since RtsA participates in a complex feed-forward loop along with HilD, we predicted that the magnitude of regulation by SdsR through the *hilD* 3′ UTR might be limited by SdsR-dependent repression of *rtsA*. To test this, we expressed SdsR in wild-type (WT) or Δ*rtsA* background strains containing the Full 3′ UTR fusion. While SdsR-dependent activation in the WT background was ∼2-fold, activation in the Δ*rtsA* background increased to ∼4-fold (Fig. 5). These data suggest that SdsR can both activate and repress SPI-1 gene expression through different targets, suggesting that this sRNA plays a role in fine-tuning SPI-1 activity through multiple independent regulatory effects.

**Figure 5.**
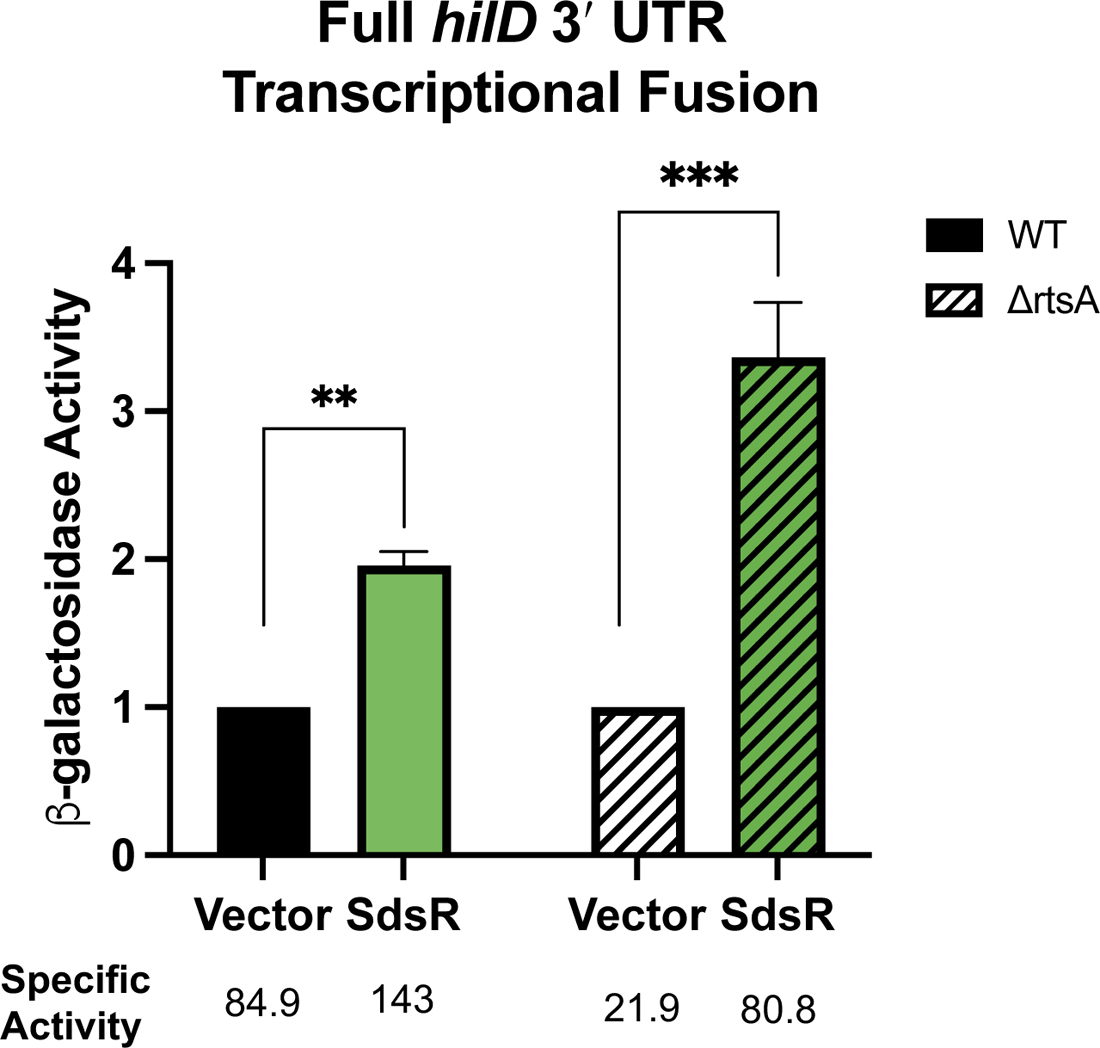
Regulation of *hilD* mRNA 3’UTR by SdsR is affected by RtsA. SdsR was produced in *Salmonella* strains containing Full *hilD* 3′UTR′-*lacZ* in a WT (*rtsA+)* or Δ*rtsA* background. Strains were grown in no salt LB, then subcultured 1:100 into high salt LB. Subcultures were grown statically for 20 hours at 37°C. Relative β-galactosidase units were obtained by normalizing β-galactosidase activity to WT control and reported as mean ± standard deviation. Error bars are standard deviations of the results of three independent experiments, obtained by unpaired *t* test, n=3. **, *P* < 0.005; ***, *P* < 0.0005, NS, not significant.

### SdsR regulates SPI-1 independently of CRP and Spot 42

Previous work on SdsR showed that one of its targets is the *crp* mRNA, which encodes the global regulator cyclic AMP receptor protein (CRP) (38). Since CRP is the transcriptional regulator that represses Spot 42 production (39), we asked whether the activation of *hilD* by SdsR could be indirect as a result of its repression of CRP (Fig. 6A). We expected that if SdsR-mediated activation of *hilD* were indirect, through repression of the repressor of Spot 42, SdsR would no longer activate the *hilD* 3′ UTR reporters in an *spf* mutant background that lacks Spot 42. To test this, we expressed SdsR and Spot 42 in in *spf+* (WT) or Δ*spf Salmonella* background strains containing the Full 3′ UTR fusion. As previously shown, in the WT background, both SdsR and Spot 42 activate the Full 3′ UTR fusion. In the Δ*spf* background, we observed ∼50% reduction in basal level of activity of the Full 3′ UTR fusion (Fig. 6B, see values for specific activity), suggesting that under these growth conditions, Spot 42 is produced and contributes positively to *hilD* expression. As expected, production of Spot 42 in the Δ*spf* background resulted in activation of *hilD.* When SdsR was expressed in the Δ*spf* strain, we saw a similar fold-induction (∼2-fold) as in the WT background (Fig. 6B), showing that SdsR acts independently of Spot 42 to activate via the *hilD* mRNA 3′ UTR.

**Figure 6:**
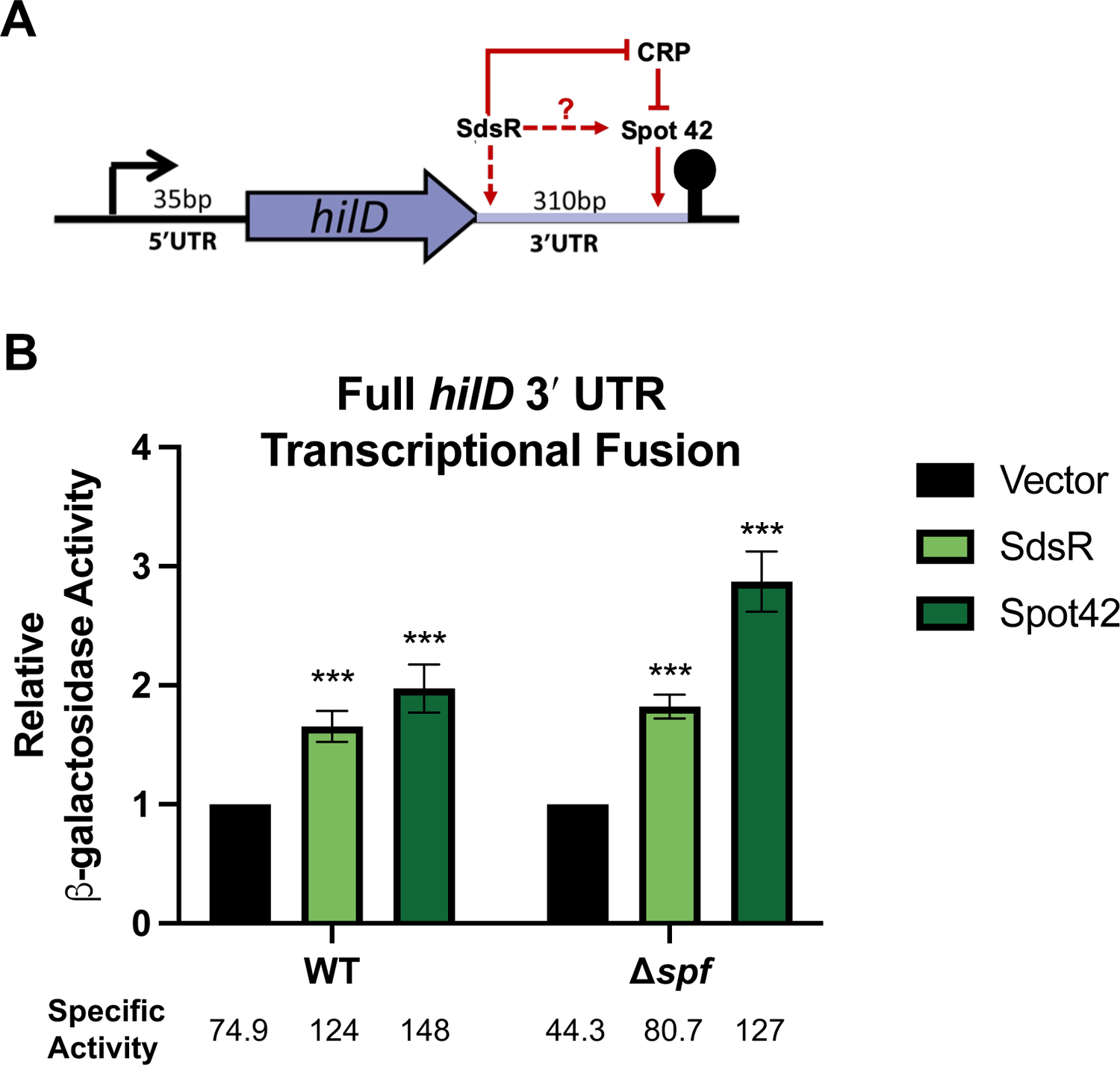
SdsR regulates *hilD* mRNA 3’UTR independently of Spot 42. (A) Model of possible indirect regulation of *hilD* mRNA 3′ UTR by SdsR and Spot 42. (B) Plasmids encoding SdsR and Spot 42 were expressed in *spf+* and *Δspf Salmonella* strains. Relative β-galactosidase units were obtained by normalizing β-galactosidase activity to WT control and reported as mean ± standard deviation. Error bars are standard deviations of the results of three independent experiments, obtained by unpaired *t* test, n=3. ***, *P* < 0.0005.

### SdsR and Spot 42 regulate *hilD* through distinct regions

Spot 42 was previously shown to interact within the region encompassing the last 185 nt of the *hilD* mRNA 3′ UTR (26). To determine whether SdsR regulates *hilD* via interactions with the same or a different region of the *hilD* mRNA 3′ UTR, we analyzed activity of SdsR and Spot 42 on the series of 3′ UTR fusions with progressive 3′ truncations (Figs. 2A, B). As observed previously (Fig. 2B and (25, 32)), activity of the fusions increased as the 3′ UTR is shortened (Fig. 7). Consistent with previous data (26), regulation of *hilD* by Spot 42 required sequences near the 3′ end of the 3′ UTR, since regulation was lost in the +180 fusion (Fig. 7). SdsR activated the Full and +180 fusions to a similar extent (∼2-fold). SdsR still activated the +90 fusion, albeit to a lesser degree than the Full and +180 fusions (Fig. 7). SdsR was not able to activate the Δ3 fusion, indicating that SdsR requires sequences within the 5′ region of the *hilD* 3′ UTR to regulate. Both SdsR and Spot 42 slightly repressed the Δ3′ UTR fusion. These data indicate that while SdsR and Spot 42 both activate *hilD* through its 3′ UTR, they target different regions to do so (Fig. 7).

**Figure 7:**
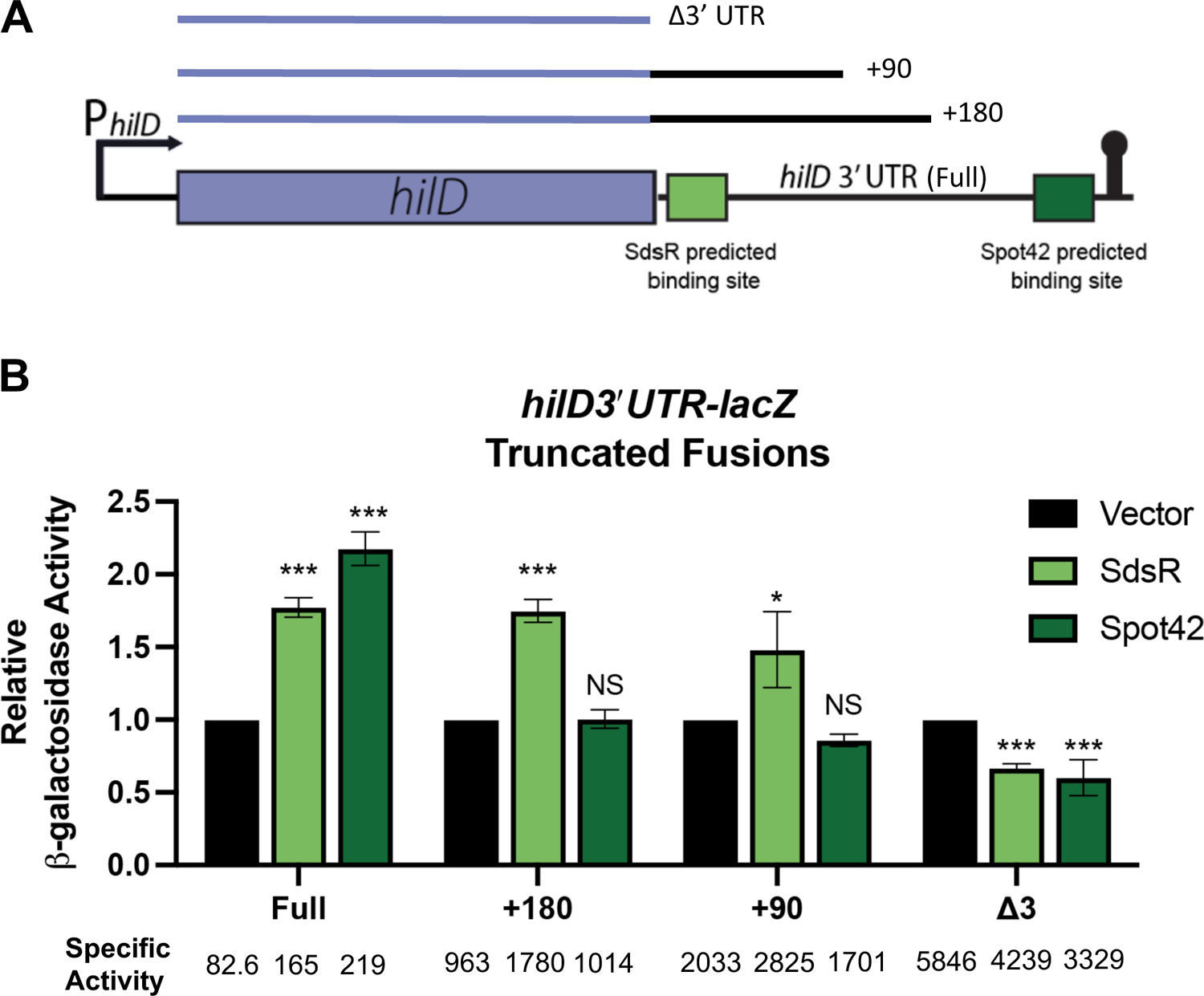
SdsR and Spot42 regulate *hilD* mRNA 3’UTR at different sites. A) Predicted base-pairing sites of sRNAs within the *hilD* mRNA 3′ UTR. B) Plasmids encoding SdsR or Spot42 (or empty vector) were expressed in strains containing the Full *hilD 3*′*UTR-lacZ*, *hilD 3*′*UTR-lacZ* +180, *hilD 3*′*UTR-lacZ* +90, or *hilDΔ3*′*UTR-lacZ* fusion constructs. Relative β-galactosidase units were obtained by normalizing β-galactosidase activity to WT control and reported as mean ± standard deviation. Error bars are standard deviations of the results of three independent experiments, obtained by unpaired *t* test, n=3. *, *P* < 0.05; ***, *P* < 0.0005, NS, not significant.

### SdsR directly interacts with *hilD* mRNA 3′ UTR

To identify specific binding interactions between SdsR and *hilD* mRNA, we used IntaRNA 2.0 to predict the binding interaction. SdsR nucleotides 47-55 (relative to the transcription start site) were predicted to base pair with *hilD* mRNA 3′ UTR nucleotides 90-98 (relative to the stop codon) (Fig. 8A). We next used RNA footprinting assays to biochemically map SdsR interactions with the *hilD* mRNA 3′ UTR. A 190-nucleotide fragment of *hilD* mRNA 3′ UTR (nucleotides 1-190 after the *hilD* stop codon) was digested with lead (II) acetate without or with SdsR and Hfq in the reactions. In the presence of SdsR alone or SdsR with Hfq, we observed an altered pattern of digestion surrounding the predicted SdsR binding site (nt 90-98), including a slight increase in protection of the putative binding site and hypersensitivity upstream and downstream from the binding site (Fig. 8B). This pattern has been observed to accompany changes in mRNA accessibility adjacent to sRNA binding sites upon sRNA binding (40–43). In the presence of both SdsR and Hfq, additional patterns of protection become evident at residues 55-58. Several A-R-N motifs, previously identified to be important for Hfq binding (44), are located upstream of this region, suggesting that this could be an Hfq interaction site. An additional protection pattern corresponding to a potential stem-loop is detected upstream between residues 39-42 and 48-51. This pattern is reproducible, and appears to represent an Hfq-induced stem-loop. Together, these results provide biochemical evidence for direct SdsR interaction with the *hilD* mRNA 3′ UTR.

**Figure 8:**
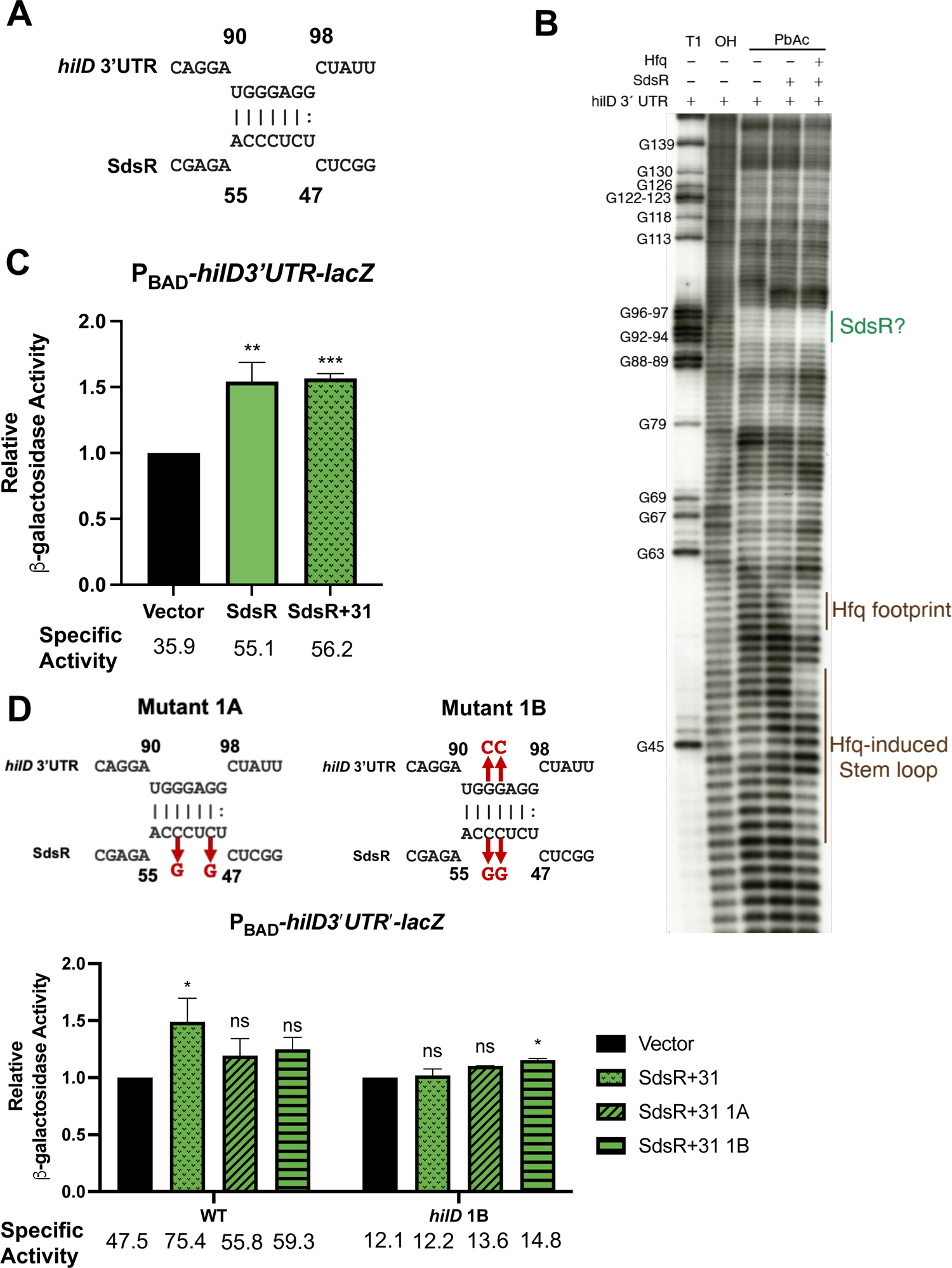
SdsR base-pairing interactions with *hilD* mRNA 3′ UTR. A) Predicted base-pairing interaction between *hilD* mRNA 3’ UTR and SdsR. B) RNA footprint of the *hilD* mRNA 3′ UTR alone, with SdsR, or with SdsR and Hfq. Nucleotide “1” corresponds to the first nucleotide downstream of the *hilD* stop codon. C) Empty vector or plasmids encoding SdsR variants were expressed in P_BAD_-*hilD 3*′*UTR-lacZ* fusion containing strain. D) SdsR variants were expressed in a strain with the P_BAD_-*hilD 3*′*UTR-lacZ* fusion, or a fusion containing the compensatory *hilD* mutations. Relative β-galactosidase units were obtained by normalizing β-galactosidase activity to vector control and reported as mean ± standard deviation. Error bars are standard deviations of the results of three independent experiments, obtained by unpaired *t* test, n=3. *, *P* < 0.05; **, *P* < 0.005; ***, *P* < 0.0005, ns, not significant.

SdsR has been previously shown to exist in two forms: full-length SdsR and a processed form called SdsR+31, which has been cleaved between nts 30 and 31, creating a new 5′ end at +31 (38). Both forms can regulate mRNA targets, depending on whether the sRNA-target interaction involves nucleotides 1-30. Since the putative interaction between SdsR and *hilD* mRNA does not involve nucleotides 1-30, we tested to see if SdsR+31 was able to regulate *hilD*. SdsR+31 regulated the full-length *hilD* 3′ UTR reporter fusion to the same degree as full-length SdsR (Fig. 8C), confirming that SdsR interaction with *hilD* mRNA does not involve the first 30nt of SdsR. To further test the SdsR-*hilD* mRNA base pairing prediction, we made mutations in the SdsR+31 construct and compensatory mutations in the *hilD* 3′ UTR reporter. SdsR variants with mutations that disrupt the putative SdsR-*hilD* mRNA interaction (SdsR+31 mutant 1A and SdsR+31 mutant 1B, Fig. 8D) fail to significantly activate the *hilD* mRNA 3′ UTR reporter fusion (Fig. 8D). A Northern blot revealed that SdsR mutant 1A was produced at lower levels than the parent SdsR+31, while mutant 1B was produced at similar levels (Fig. S4), suggesting that loss of regulation of *hilD* by SdsR mutant 1B was due to disruption of base pairing. To test if regulation could be restored by compensatory changes in the *hilD* 3′ UTR, we created a P_BAD_-*hilD3′UTR-′lacZ* fusion containing the mutations that should restore base pairing with SdsR mutant 1B (*hilD* 1B, Fig. 8D). Consistent with the base pairing prediction, wild-type SdsR+31 no longer regulated the *hilD* 1B reporter fusion (Fig. 8D), but the regulation by SdsR mutant 1B was not fully restored. Nevertheless, loss of regulation caused by mutations in either SdsR or *hilD* mRNA binding sites provides genetic evidence consistent with the biochemical evidence (Fig. 8B) supporting our model of SdsR-mediated regulation of *hilD* via a direct base pairing interaction between SdsR and the *hilD* mRNA 3′ UTR.

### SdsR regulates SPI-1 in an RNase E-dependent manner

sRNAs often regulate mRNA processing or stability by modulating RNase E access to or cleavage activity on mRNAs. The *hilD* mRNA 3′ UTR has been proposed to contain numerous RNase E cleavage sites (25, 34), and we have shown that *hilD* 3′ UTR-mediated regulation is dependent on RNase E (Fig. 3). We propose that SdsR may stabilize the *hilD* mRNA 3′ UTR by occluding RNase E cleavage sites. To determine if SdsR-mediated regulation of *hilD* involves RNase E, we compared activity of SdsR on the full-length *hilD* 3’UTR reporter fusion in wild-type and RNase E degradosome mutant (*rne131*) backgrounds. The basal level of activity of the reporter fusion increased 4-fold in the *rne131* background compared to the wild-type background (Fig. 9A, see specific activity values below the graph), consistent with the model that the *hilD* 3′ UTR is a target for RNase E. While SdsR and Spot 42 both activated the *hilD* 3′ UTR reporter fusion in a wild-type (*rne*^+^) background, SdsR lost the ability to activate in the *rne131* background, while Spot 42-mediated regulation was impaired (Fig. 9A).

**Figure 9:**
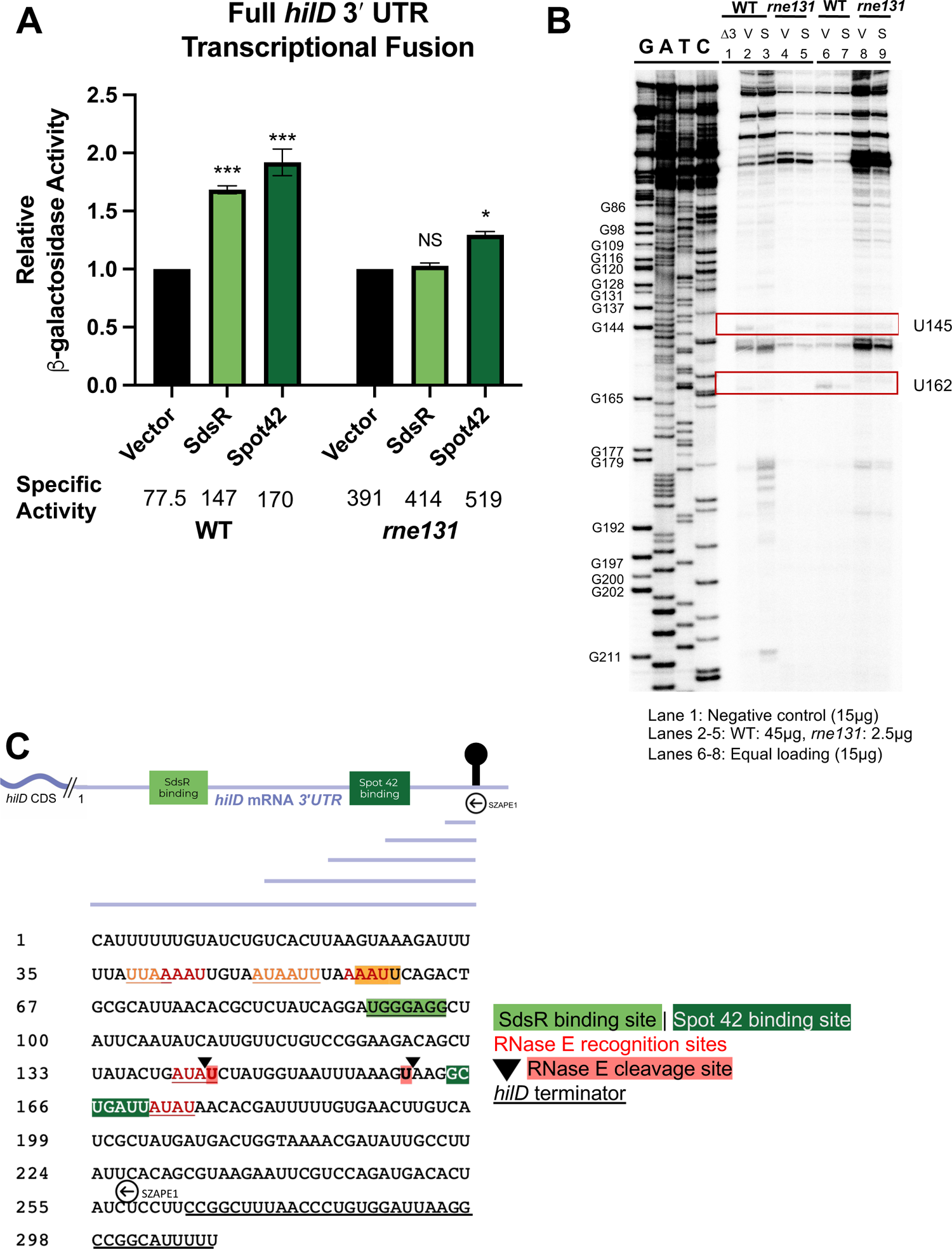
RNase E plays a role in sRNA regulation of the *hilD* mRNA 3’UTR. (A) SdsR and Spot 42 were produced in *Salmonella* strains containing the Full *hilD* 3′UTR′-*lacZ* fusion in a WT or *rne131* background. Relative β-galactosidase units were obtained by normalizing β-galactosidase activity to vector control and reported as mean ± standard deviation. Error bars are standard deviations of the results of three independent experiments, obtained by unpaired *t* test, n=3. *, *P* < 0.05; **, *P* < 0.005; ***, *P* < 0.0005, NS, not significant. (B) Primer extension assay of *hilD* mRNA 3′ UTR using primer SZAPE1 in the presence or absence of SdsR in either a wild-type or *rne131* background. V=vector control; S=SdsR. Sanger ladder created with SZAPE1 template, nucleotide “1” corresponds to first nucleotide downstream of *hilD* stop codon. Lane 1: RNA harvested from Δ3’UTR strain (15µg), Lanes 2-5: Loaded to provide comparable band intensity (45µg WT samples, 2.5µg *rne131* samples), Lanes 6-9: Equal loading of RNA from WT vs *rne131* samples (15µg). The red box indicates products whose abundance changes in an SdsR-dependent manner. (C) Diagram of *hilD* 3’ UTR and expected primer extension products. Nucleotide sequence of *hilD* mRNA 3’UTR including sRNA binding sites, RNase E recognition sites (34) and relevant ends from primer extension. Red highlight corresponds to red box in (B), red text corresponds to predicted RNase E recognition sites (34).

We next wanted to map cleavage sites within the *hilD* 3′ UTR by using primer extension to identify cleaved transcripts in wild-type (*rne*+) and *rne131* backgrounds and compare whether there were differences in the presence of SdsR. Total RNA was extracted from *rne*+ and *rne131* strains containing the vector control or SdsR expression plasmids. A primer situated just upstream of the *hilD* 3’ terminator directed synthesis towards the 5′ of the *hilD* transcript. Shorter fragments running toward the bottom of the gel presumably represent cleavage of the mRNA closer to the terminator, whereas longer fragments toward the top of the gel represent cleavage closer to the *hilD* stop codon (Fig. 9B, 9C). As expected, there was no reaction for the negative control sample (τ<*hilD3′ UTR*, Lane 1, Fig. 9B). Because the abundance of *hilD* mRNA increases significantly in the *rne131* background (Fig. 9A), when equal amounts of RNA were loaded for wild-type and *rne131* samples (15 μg, Fig. 9B, lanes 6-9), the intensity of bands in the wild-type samples was too weak to identify differences between samples. Nevertheless, an overall comparison between the wild-type samples (lanes 6, 7) and samples from the *rne131* strain (lanes 8, 9) revealed a substantial accumulation of *hilD* 3′ UTR species of various sizes, including longer species.

To better compare the patterns between wild-type and *rne131* samples, in lanes 2-5 (Fig. 9B) more RNA was loaded for wild-type samples (45 μg) than for *rne131* samples (2.5 μg) to achieve a comparable intensity of bands between the samples. Comparing wild-type (*rne*+) samples without (lane 2) and with (lane 3) SdsR, we observed increased levels of certain products in the presence of SdsR, suggesting that SdsR binding changed susceptibility to cleavage at one or more sites in the *hilD* 3′ UTR. There was no noticeable difference between samples in the *rne131* strain without (lane 4) or with (lane 5) SdsR, consistent with the above interpretation.

If SdsR suppressed cleavage at specific sites in the *hilD* 3′ UTR, we would expect to see reduced intensity of specific bands in the wild-type samples with SdsR compared to without SdsR. We identified two such bands. First, a band corresponding to a cleavage at residue A145 was present without SdsR (Fig. 9B, lane 2) and was decreased in abundance in samples where SdsR was expressed (lane 3). Interestingly, this region corresponds to one of the putative RNase E processing sites predicted by Song et al. (34), and is downstream of the putative SdsR binding site, suggesting a possible role for SdsR-mediated occlusion of an RNase E processing site, perhaps through a structural shift. A second band corresponding to residue A162 was also decreased in abundance when SdsR was expressed, and this effect was also seen in an *rne131* background. Interestingly, this region is near the putative Spot 42 binding site. Comparing the digestion pattern of the RNA footprint with these residues of interest, A145 appears to be in a single-stranded region that would be accessible to RNase E, and the region appears to have slightly increased protection upon SdsR binding (Fig. S5). Understanding the identity and role of other *hilD* 3′ UTR regulators, as well as more in-depth structural analysis of the *hilD* mRNA 3′ UTR, will provide more insight. Combined, these observations provide preliminary evidence that structural shifts may be playing a role in occluding RNase E processing sites and protecting the *hilD* transcript.

### SdsR and Spot 42 affect *Salmonella* virulence in mice

SdsR and Spot 42 act through the *hilD* mRNA 3′ UTR to increase SPI-1 expression. We wanted to examine if this regulation impacts infection. To understand the importance of SdsR and Spot 42 during infection, we performed mouse competition assays, recovering strains from the lower small intestine and the spleen. We have previously demonstrated that decreased recovery of a strain from the spleen after oral infection is consistent with defects in invasion (8). Intraperitoneal (i.p.) competition assays are also performed with the expectation that SPI-1 dependent defects will not be reflected in spleens after i.p. infection, which bypasses the need for SPI-1-mediated invasion.

After oral infection, recovery of the Δ*sdsR* Δ*spf* strain was significantly reduced in the spleen (Table 1). Recovery of the Δ*sdsR* Δ*spf* from the small intestine was also decreased, but results were not statistically significant (Table 1). There was no defect observed in Δ*sdsR* Δ*spf* strains recovered from spleens after i.p. infection (Table 1), suggesting strongly that the phenotype observed after oral infection reflects defects in intestinal colonization and/or invasion.

**Table 1:** SdsR and Spot 42 affect intestinal colonization and invasion.

| Table 1: SdsR and Spot42 affect intestinal colonization and invasion |  |  |  |  |  |  |
| --- | --- | --- | --- | --- | --- | --- |
| Strain A<sup>a</sup> | Strain B<sup>a</sup> | Infection route<sup>b</sup> | Organ<sup>b</sup> | No. of mice | Median CI<sup>c</sup> | P value<sup>d</sup> |
| $\Delta sdsR\Delta spot42$ | WT | Oral | Spleen | 10 | 0.21 | 0.02 |
|  |  | Oral | SI | 10 | 0.52 | NS |
|  |  | i.p | Spleen | 10 | 1.83 | 0.04 |
| $\Delta sdsR\Delta spot42$<br>$\Delta spi-1$ | $\Delta spi-1$ | Oral | Spleen | 10 | 8.25 | NS |
|  |  | Oral | SI | 10 | 10.5 | 0.01 |
|  |  | i.p | Spleen | 9 | 1.46 | NS |
| $\Delta spi-1$ | WT | Oral | Spleen | 5 | 0.09 | <0.005 |
|  |  | Oral | SI | 5 | 0.03 | 0.01 |
|  |  | i.p | Spleen | 5 | 0.98 | NS |
| $\Delta sdsR\Delta spot42$<br>$\Delta spi-1$ | $\Delta sdsR$<br>$\Delta spot42$ | Oral | Spleen | 10 | 0.25 | 0.03 |
|  |  | Oral | SI | 10 | 0.31 | 0.03 |
|  |  | i.p | Spleen | 7 | 0.95 | NS |

NS<sup>e</sup> NS<sup>e</sup>
<sup>a</sup>The strains used were JS135, SZA422, 423, SZA433-436 (Table S1).
<sup>b</sup>Bacteria were recovered from the spleen (Sp) after intraperitoneal (i.p.) infections or from the small intestine (SI) and spleen after oral infection.
<sup>c</sup>The competitive index (CI) was calculated as (percentage of strain A recovered/percentage of strain B recovered)/(percentage of strain A inoculated/percentage of strain B inoculated).
<sup>d</sup>Student's *t* test was used to compare the CIs to the inocula, or between groups as indicated. ns, not significant (*P*>0.05).
<sup>e</sup> Student's *t* test was used to compare oral infection groups " $\Delta spi-1$ vs WT" and " $\Delta sdsR\Delta spot42\Delta spi-1$ vs $sdsR\Delta spot42$ "

To determine if this observed defect is dependent on SPI-1, we performed oral and i.p. infections in a Δ*spi-1* background. The Δ*sdsR* Δ*spf* Δ*spi-1* strain was not attenuated relative to the Δ*spi-1* strain. Indeed, Δ*sdsR* Δ*spf* Δ*spi-1* strain significantly outcompeted the Δ*spi-1 sdsR+*/*spf*+ strain in the small intestine and was recovered in relatively higher numbers from spleens after oral infection (Table 1). There was no significant difference in recovery of the two strains after i.p. infection. These results were reproducible; these strains were independently reconstructed, and the infections repeated, obtaining essentially equivalent results. The data in Table 1 are combined from the independent experiments. While the reason for this phenomenon is unclear, both SdsR and Spot 42 have a broad spectrum of targets and are apparently affecting some aspect of *Salmonella* physiology that allows increased colonization or growth in the small intestine, resulting in increased numbers of bacteria crossing the intestinal epithelium via a SPI-1-independent mechanism (45).

The simplest explanation for the results above is that loss of SdsR and Spot42 confers two opposing phenotypes: decreased expression of SPI-1 with the expected loss in invasion, and some change in physiology that increases colonization, independent of SPI-1. The SPI-1 defect is dominant. To further test this model, we determined the effect of deleting SPI1 in the Δ*sdsR* Δ*spf* background, with the expectation that, since SPI1 expression was already reduced, deleting the locus would have less of an effect compared to deleting SPI-1 in the wild-type background. Indeed, we observed a decreased defect compared to that observed in the Δ*spi-1* vs WT oral competition. Although there is no statistical significance between the groups, the trend corresponds to our observations in previous assays. No effect was observed after i.p. infection consistent with all the effects being limited to the intestine. Overall, results confirm that the initial defect observed in WT oral infection is SPI-1 dependent. While further studies are required to understand how the sRNAs affect intestinal colonization, we can conclude that SdsR and Spot 42 play a role in facilitating SPI-1 mediated invasion.

## Discussion

*Salmonella* replicates within various niches in the host (46), sensing a variety of environmental cues to tune virulence gene expression, including the genes for invasion, encoded on *Salmonella* pathogenicity island 1 (SPI-1). The SPI-1 genes must be tightly regulated in space and time, as too little or too much expression of the SPI-1 Type 3 secretion system impairs virulence (7). We have previously identified multiple sRNA regulators that post-transcriptionally regulate SPI-1 through interactions with the 5′ UTRs of the *hilD* and *hilA* mRNAs (27–29) to help maintain this tight level of regulation. The *hilD* 3′ UTR represents another target site for sRNA regulators, and it was previously shown that the sRNA Spot 42 acts as a positive regulator of SPI-1 through interactions with the *hilD* 3′ UTR (25, 26). In this study, we identified an additional sRNA, SdsR, that activates SPI-1 via the *hilD* mRNA 3′ UTR (Figs. 2, 4). While SdsR and Spot 42 activities are connected through the transcriptional regulator CRP, we showed that SdsR regulates *hilD* independent of Spot 42 (Fig. 6) and SdsR and Spot 42 target different regions of the *hilD* mRNA 3′ UTR (Fig. 7). We provide evidence that SdsR positively regulates *hilD* by altering susceptibility of the 3′ UTR to cleavage by RNase E (Fig. 9). Collectively, our results support a model in which the *hilD* 3′ UTR makes *hilD* mRNA susceptible to RNase E-dependent degradation, and the sRNAs Spot 42 and SdsR protect the mRNA from cleavage by RNase E, yielding higher levels of HilD and higher SPI-1 gene expression (Fig. 1,10). We have found that while both SdsR and Spot 42 activate *hilD*, they target *hilD* differently – SdsR targets a binding site within the first 90 nt of the *hilD* 3′ UTR while Spot 42 targets a region closer to the terminator. Consistent with other studies (20, 25, 34), these observations suggest that Hfq and RNase E regulate mRNA stability by acting at multiple sites across the long *hilD* 3′ UTR.

**Figure 10.**
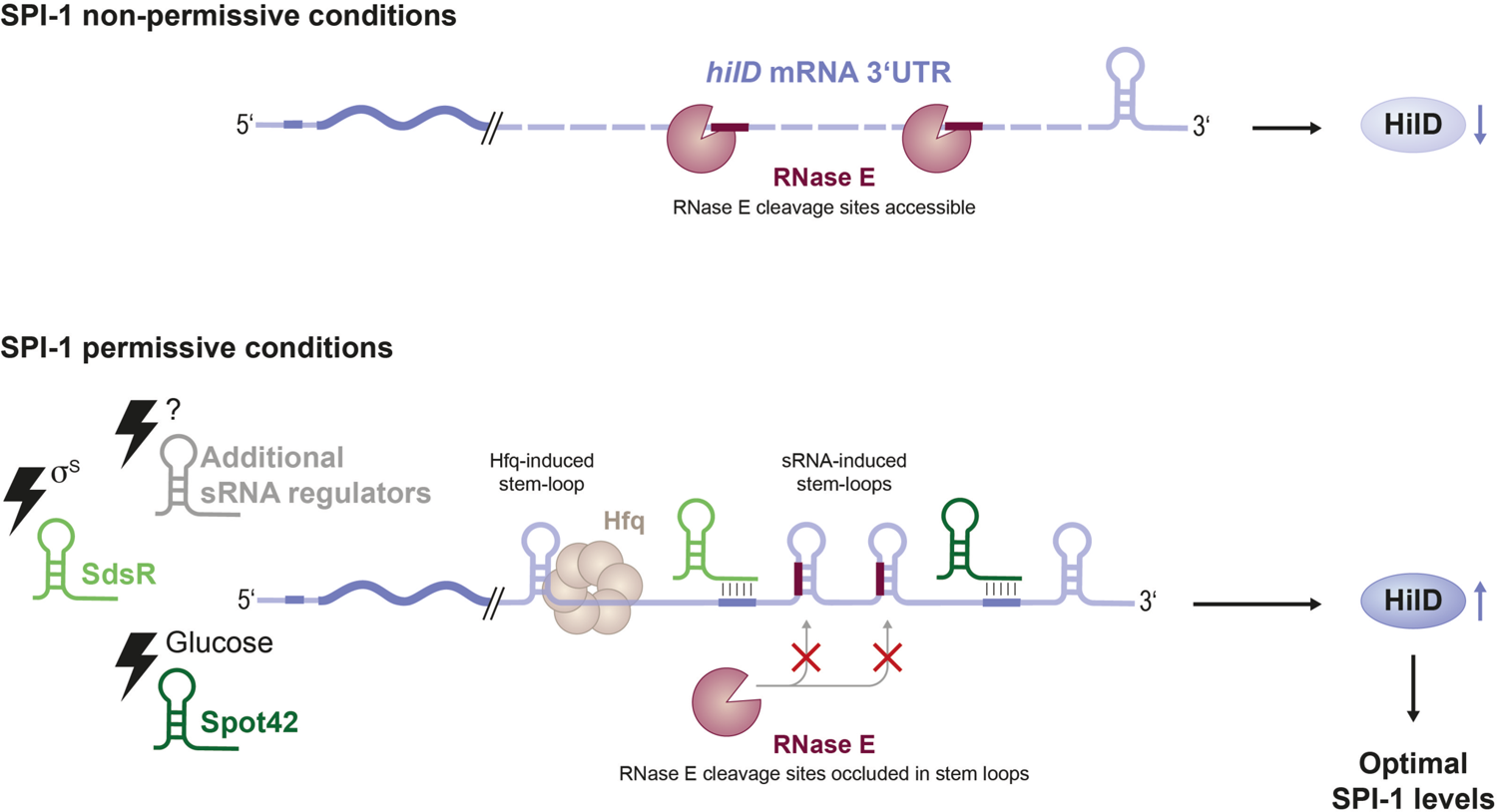
sRNA mediated regulation of SPI-1. Working model for the SdsR- and Spot 42-mediated regulation of SPI-1. The *hilD* 3**′** UTR contains multiple RNase E cleavage sites, making *hilD* mRNA inherently unstable under conditions where SPI-1 is not induced. A variety of conditions associated with SPI-1 induction can induce production of sRNAs such as Spot 42 and SdsR. Along with Hfq, sRNAs can base pair with *hilD* mRNA 3**′** UTR to promote structural rearrangements that occlude RNase E sites and protect *hilD* mRNA from RNase E-mediated degradation.

The regulation of *hilD* by sRNAs targeting the 3′ UTR represent some of the first reports of regulation of 3′ untranslated regions by bacterial small RNAs. While eukaryotic microRNAs (miRNAs) often target 3′ UTRs, the regulatory roles of bacterial 3′ UTRs are not well characterized, aside from a few recent discoveries. The *Staphylococcus aureus* biofilm repressor, IcaR possesses a long 3′ UTR that acts in *cis* via an anti-Shine Dalgarno (SD) sequence that inhibits translation via intramolecular base pairing (47). Bronesky, *et al.,* showed that the sRNA, RsaI, regulates *icaR* mRNA through interactions with its 3′ UTR to disrupt this intramolecular interaction (48). The *hilD* mRNA 3′ UTR similarly acts as a regulatory element, but in this case as an independent regulatory module capable of conferring regulation at the *hilD* locus, or at an exogenous locus (Fig. 2D, Fig S2 (25, 34)). This regulatory module limits expression of HilD, and due to the SPI-1 complex feed-forward loop (Fig. 1), in turn, the signal is amplified to maintain tight repression of SPI-1 genes (Fig. 1, 10).

While details of regulation at 3′ UTRs are still being explored, a feasible mechanism for sRNA-mediated regulation involves directly occluding RNase E recognition sites or remodeling the structure to reveal or hide RNase E recognition sites. Previous studies have determined that RNase E plays a role at the *hilD* mRNA 3′ UTR to maintain low levels of *hilD* mRNA (25). We found that in an *rne131* background, certain *hilD* mRNA 3′ UTR-derived fragments were more abundant (Fig. 9B), suggesting that an intact RNA degradosome is required to promote rapid and complete degradation of this region of the *hilD* mRNA. When SdsR was expressed in a wild-type (*rne*^+^) background, we observed changes in accumulation of certain *hilD* 3′ UTR-derived fragments, including fragments that are not observed in an *rne131* background, consistent with SdsR altering susceptibility of the 3′ UTR to RNase E-dependent processing.

We identified two potential RNA processing sites based on the decreased abundance of fragments in the presence of SdsR, or in an *rne131* background. One such site, involving nucleotide U145, relative to the *hilD* stop codon, appears to be in a single-stranded region that has slight protection upon SdsR binding (Fig. 9, Sig. S5), consistent with our hypothesis that sRNA binding is protecting *hilD* mRNA by occluding RNase E processing sites within stem-loops. The second site, involving U162 appears to be in an already protected region, and is located close to the putative Spot 42 binding site (Figure 9B, (26)), but could potentially become accessible based on the overall shift in structure that occurs upon sRNA binding. Both sites are located downstream of the putative SdsR binding site (Fig. 9C, 10). Although this could suggest multiple binding sites for SdsR, we did not identify any promising putative binding interactions in this region. Instead, we propose that these findings, combined with shifts in hypersensitivity observed in the RNA footprinting, provide evidence for SdsR-induced changes in *hilD* mRNA 3′ UTR structure. Further structural studies could provide the necessary insights to understand the dynamics at the *hilD* mRNA 3′ UTR.

SdsR is an RpoS-dependent sRNA with a large target regulon (38). Among its targets are mRNAs encoding transcriptional regulators such as CRP and RtsA, which adds additional layers of complexity to the SdsR network. SdsR negatively regulates CRP, which is the transcription factor controlling Spot 42 production. Our work shows that SdsR regulates *hilD* independently of Spot 42 (Fig. 6B), suggesting that regulation of *hilD* and *crp* by SdsR may occur under different conditions, or may work together to allow maximal SPI-1 activation at the correct time. SdsR regulation of *rtsA* mRNA (27) is of particular interest, as RtsA and HilD are part of the same regulatory loop that activates expression of SPI-1 (Fig. 1). We have previously shown that the sRNA PinT also regulates both *hilD* and *rtsA*, but in this case, PinT represses both (27). Here, we find that SdsR represses *rtsA* while upregulating *hilD* (Fig. 5). We have previously demonstrated that SPI-1 regulators form homodimers and heterodimers (49). The various forms of heterodimer will differ depending on the relative concentrations of the regulators and different homodimers or heterodimers likely differ in their ability to bind to or activate certain promoters (49). Differential regulation of RtsA and HilD by SdsR would shift the relative concentrations of heterodimer and homodimers, consistent with sRNAs acting to fine-tune precise levels of gene expression rather than as simple ON/OFF switches.

We have shown that SdsR and Spot 42 play a role in facilitating *Salmonella* infection (Table 1). The predominant effect is a decrease in SPI-1-mediated invasion, as would be expected from decreased HilD production in the *sdsR spf* mutant background. Interestingly, in the absence of SPI-1, deletion of SdsR and Spot 42 seems to provide a competitive advantage during oral infection (Table 1), suggesting that these sRNAs are somehow hindering SPI-1-independent invasion. It has previously been shown that *Δspi-1* strains are capable of invading into deeper tissue via CD-18 expressing phagocytes (45). Both SdsR and Spot 42 have numerous targets that could directly or indirectly mediate this effect. The ability of this mutant to outcompete an *sdsR^+^spf*^+^ strain in a *Δspi-1* background provides further evidence of the intricacies of regulatory networks and cross-talk surrounding SPI-1 infection.

We propose that SdsR and Spot 42, along with Hfq, activate *hilD* through different regions of the *hilD* mRNA 3′ UTR, and sRNA binding protects *hilD* mRNA from RNase E-mediated degradation under SPI-1 permissive conditions, allowing for optimal HilD levels. We have identified putative sites of protection downstream of the SdsR binding site and propose that this protection occurs through shifts in the RNA structure, potentially occluding single-stranded RNase E processing sites in structured loops (Fig. 10). While we have preliminary evidence to support this, further structural studies of the 3′ UTR are necessary to provide more insight and confirm these observations. In addition to our findings, the role of Gre factors at the *hilD* 3′ UTR suggests a role for RNA polymerase pausing or backtracking (32), which could also aid in modulating different structures or in allowing for a sRNA-stabilized form of *hilD* 3′ UTR during permissive conditions. In modulating mRNA stability and potentially causing structural shifts to do so, bacterial sRNAs may play similar roles to eukaryotic miRNAs (50, 51). Many questions remain about the role of other *hilD* 3′ UTR regulators and how they interact with one another to regulate HilD levels. This work gives us a glimpse into how sRNAs regulate at 3′ UTRs, thus opening the door to better understanding the roles of 3′ UTRs in bacteria, as well as providing additional details of SPI-1 regulation.

## Materials and Methods

### Strain and plasmid construction

Strains and plasmids used in this study are listed in Table S1. All *Salmonella enterica* serovar Typhimurium strains used in this study are derivatives of ATCC [American Type Culture Collection] 14028s. Chromosomal mutations were made using λ red recombination (52) and moved into the appropriate strain background using P22 HT-105/1 *int-*201-mediated transduction (53).

Transcriptional *lacZ* fusions in *E. coli* were constructed by λ red recombination in the strain PM1805 as described previously (35). Resulting fusions are under the control of arabinose-inducible P_BAD_ promoter. All oligonucleotide primers used in this study were synthesized by Integrated DNA Technologies and are listed in Table S2. PCR products were generated using Q5 Hot Start High-fidelity system (New England Biolabs). Plasmids encoding IPTG-inducible *sdsR* or *spf* (Spot 42) were constructed by PCR amplification of *sdsR* and *spf* genes from strain 14028s using primers F-SdsR_aatII/R-SdsR_ecoRI and F-spf_aatII/R-spf_ecoRI, respectively (Table S2). The PCR products were cloned into a pBR-P_Lac_ vector digested with *Aat*II and *Eco*RI (New England Biolabs)(54) and dephosphorylated with calf intestinal alkaline phosphatase (New England Biolabs). gBlocks (Integrated DNA Technologies) were used to construct plasmids containing the SdsR+31 mutants, with gBlocks cloned into pBR-P_Lac_ following restriction digestion with *Aat*II and *Eco*RI. gBlocks containing the *hilD* 3′ UTR were used to construct the P_BAD_-*hilD 3′UTR′-lacZ* fusions and mutant variants in the strain PM1805 using λ Red recombination (35). IntaRNA 2.0 was used to obtain predictions for base pairing between SdsR and *hilD* mRNA 3’ UTR (55).

### Media and Growth Conditions

Bacterial strains were cultured in LB medium containing 10 g tryptone and 5 g yeast extract per liter (no salt LB, NSLB), or 10 g tryptone, 5 g yeast extract, and 10 g NaCl per liter (high salt LB, HSLB) at 37°C as indicated in figure legends. Strains harboring temperature-sensitive plasmids such as pSIM6, pKD46, and PCP20 were cultured at 30°C. When necessary, antibiotics were added at the following concentrations: 100 μg/mL ampicillin (amp), 25 μg/mL kanamycin (kan), 25 μg/mL chloramphenicol (cm), 20 μg/mL tetracycline (tet), 50 μg/mL apramycin (apr). Bacteria were cultured in NSLB, or HSLB broth, or on LB plates at 37°C. Where appropriate, 0.1 M Isopropyl-β-D-1-thiogalactopyranoside (IPTG) was used for induction of the P_lac_ promoter.

### β-galactosidase Assays

*Salmonella* strains containing *lacZ* reporter fusions were cultured overnight in NSLB with the appropriate antibiotics, then subcultured 1:100 into HSLB and grown at 37°C statically for 20 hours (SPI-1 inducing conditions). β-galactosidase assays were performed using a kinetic microtiter plate assay described previously (56) using the conditions indicated in the figure legend. β-galactosidase units are defined as μmol of ortho-nitrophenyl-β-galactoside formed min^-1^ X 10^6^/(OD_600_ x mL OD cell suspension) and are reported as mean ± standard deviation of three independent experiments (n=3), analyzed statistically using an unpaired student *t* test.

*E. coli* strains containing *lacZ* reporter fusions were cultured overnight in LB with the appropriate antibiotics, then subcultured 1:100 into fresh LB medium containing 0.002% L-arabinose. Strains were grown at 37°C with shaking to early exponential phase. Samples testing ectopic sRNA expression were induced with 100 μM IPTG (to induce sRNA expression) for 1 hour, harvested, and β-galactosidase assays were performed as described above.

### *In vitro* transcription

Template DNA for in vitro transcription was produced by PCR using oligonucleotides incorporating the T7 promoter sequence on the 5′ end of the forward primer. Primers specific to the *hilD* mRNA 3′ UTR (o-D3ShT7F, o-hilD3R) and SdsR (o-T7sdsRF, o-sdsRR) were used to amplify *hilD* 3′ UTR and SdsR, respectively, from genomic DNA of *Salmonella* 14028s. *In vitro* transcription was performed using MEGAscript T7 kit (Ambion) according to the manufacturer’s instructions.

### RNA Footprinting

RNA footprinting reactions were performed as described previously (57). In short, *in vitro* transcribed *hilD* mRNA was labelled using KinaseMax (Ambion) according to the manufacturer’s instructions. 0.1 pmol of 5′-end labeled *hilD* 3′ UTR mRNA was incubated with 100 pmol of unlabeled SdsR for 10 minutes in structure buffer (Ambion) and 1 ng of yeast RNA (Ambion). 2.5 uM lead acetate was added for 2 minutes at 37 °C. 12 μL of loading buffer II (Ambion) was added to stop the reactions. A size ladder created by alkaline hydrolysis of RNA was generated by boiling the *hilD* 3′ UTR transcript at 90°C in alkaline buffer. A G-ladder was generated by incubating *hilD* 3′ UTR and RNase T1 for 5 minutes at 37°C. Samples were resolved on an 8% polyacrylamide-urea gel and imaged using GE Typhoon FLA 7000 phosphor imaging software.

### RNA Isolation and Northern Blot Analyses

Total RNA was isolated using the hot phenol RNA extraction method, as previously described (58). Between 1 and 5 μg of RNA was boiled at 95 °C with Gel II Loading Buffer (Invitrogen) and loaded on a urea-PAGE gel (Sequagel, National Diagnostics). Samples were run for 1-2 hours at 100V in 1X TBE buffer, transferred onto BrightStar™-Plus Positively Charged Nylon Membrane (Ambion) in 0.5X TBE or Northern MAX transfer buffer (Ambion), and cross-linked by 0.12 J/cm2 UV light. Oligonucleotide probes from IDT hybridizing to SdsR or 5S rRNA were end-labelled using KinaseMax 5′ end-labelling kit (Ambion). Blots were hybridized overnight with ^32^P-5′ end-labelled probes at 42°C, washed twice with wash buffer 1 (2X SSC–0.1% SDS) for 15 minutes, then washed twice with wash buffer 2 (0.1X SSC–0.1% SDS) for 15 minutes. Gels were imaged using GE Typhoon FLA 7000 phosphor imaging software.

### Primer Extension

Primer extension was performed as previously described (59). Briefly, total RNA was isolated using the hot phenol RNA extraction method, as previously described (58). Total RNA was converted to cDNA using SuperScript IV-RT (Invitrogen). An oligonucleotide corresponding to *hilD* mRNA 3′ UTR (PE1-hilD) that was 5′-end-labelled with γ-^32^P was used for reverse transcription of total RNA at concentrations specified in figure legends. A ladder was generated with Sequenase 2.0 (Affymetrix) using oligonucleotides PE1-hilD and PE3-hilD. Samples were prepared with Gel Loading Buffer II (Invitrogen), boiled for 2 minutes, then run on an 8% Urea-polyacrylamide gel (SequaGel, National Diagnostics) at 40W for 2.5 hours. Gels were imaged using GE Typhoon FLA 7000 phosphor imaging software.

### Animal Infections

BALB/c mice (Envigo) (10 to 13 weeks old) were inoculated either orally or intraperitoneally (i.p.) with 0.2 ml of a bacterial suspension of a 1:1 ratio of two bacterial strains. Each inoculum was plated on LB agar medium to measure the total inoculum and replica plated to the appropriate selective medium to calculate the input ratio for each strain. Bacterial suspensions were prepared from 16-hour overnight cultures in NSLB, diluted to the appropriate concentration in oral saline (pH 8.0) or 1x PBS. For oral infections, the bacteria were washed and suspended at 5 × 10^8^ CFU (wild-type background) or 10^9^ CFU (*Δspi1* background) per 0.2 ml. Food and water were withheld 4 hours prior to oral infection and replaced immediately after infection. For intraperitoneal infections, cells were diluted to 10^3^ CFU per 0.2 ml in sterile phosphate-buffered saline (1x PBS). Mice were sacrificed by CO_2_ asphyxiation at 3.5 days after oral infection, or 4.5 days after i. p infection. Organs were homogenized, and serial dilutions were plated on LB medium and replica plated to appropriate selective medium to calculate the output ratio for each competition. The competitive index (CI) was calculated as (percent strain A recovered/percent strain B recovered)/(percent strain A inoculated/percent strain B inoculated).. Student’s t-test was used to determine whether the output ratio was significantly different from the input ratio, and for comparison between groups. Data from *ΔΔsdsRspf* strain is a compilation of two independently built strains containing the *ΔΔsdsRspf* deletion.

## Ethics statement

All animal work was reviewed and approved by the University of Illinois Institutional Animal Care and Use Committee (IACUC). Procedures were performed in the AAALAC-accredited facility in accordance with University and PHS guidelines under protocol 21197. All efforts were made to minimize animal suffering.

## Acknowledgements

This work was supported by National Institutes of Health grant R01 GM120182 to C.K.V. and J.M.S., R21 AI166495 to J.M.S., and R35 GM139557 to C.K.V., as well as a University of Illinois Department of Microbiology Marie Chow Teaching Scholarship and Francis M. and Harlie M. Clark Microbiology Fellowship to S.Z.A. We thank Sandy Pernitzsch from SciGraphix for graphic design of model figures. Finally, we thank the current and former members of the Vanderpool and Slauch laboratories for strains, plasmids, advice, and helpful discussions.

---

**Figure S1:**
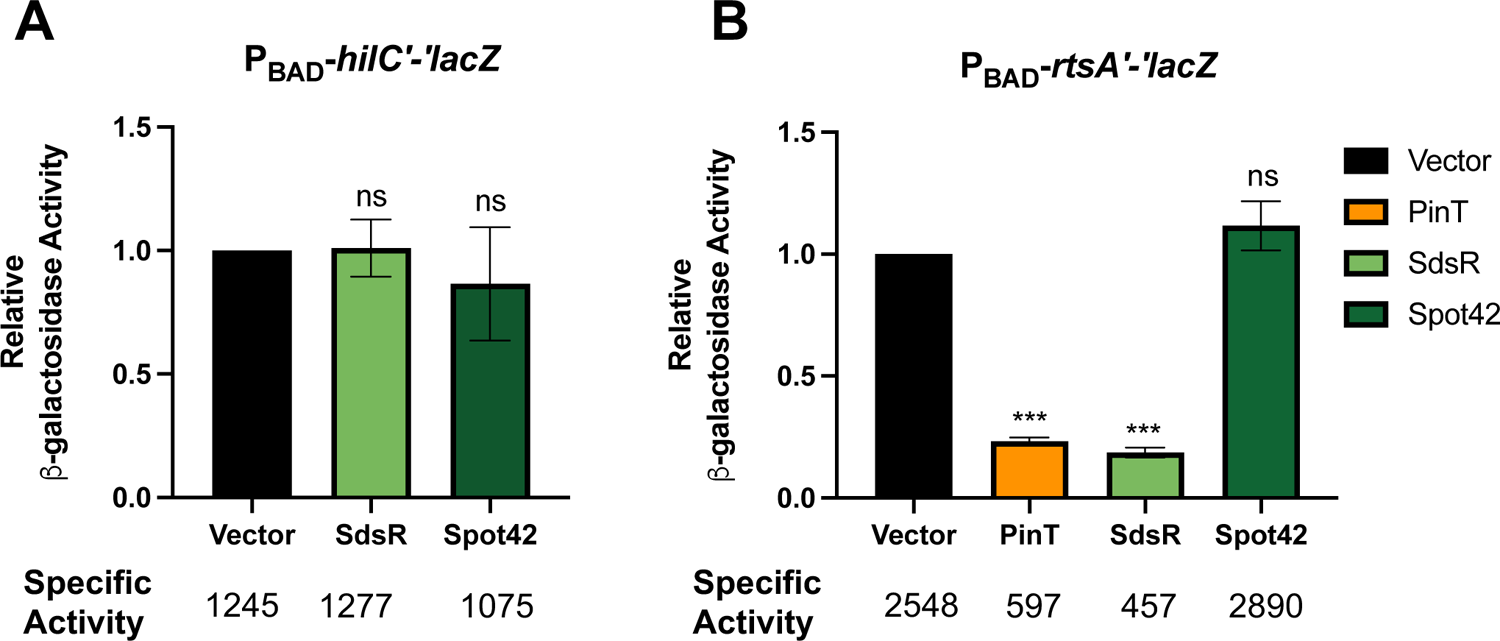
SdsR regulates RtsA but not HilC. SdsR and Spot 42 were expressed in P_BAD_-*hilC*′*-*′*lacZ* (A) or P_BAD_-*rtsA*′*-*′*lacZ* (B) translational fusions to see whether they regulate HilC or RtsA expression. A) SdsR and Spot 42 do not regulate *hilC.* B) SdsR represses RtsA. PinT represses RtsA and serves as a positive control. Spot42 does not affect RtsA. Relative β-galactosidase units were obtained by normalizing β-galactosidase activity to WT control and reported as mean ± standard deviation. Error bars are standard deviations of the results of three independent experiments, obtained by unpaired *t* test, n=3. ***, *P* < 0.0005, ns, not significant.

**Figure S2:**
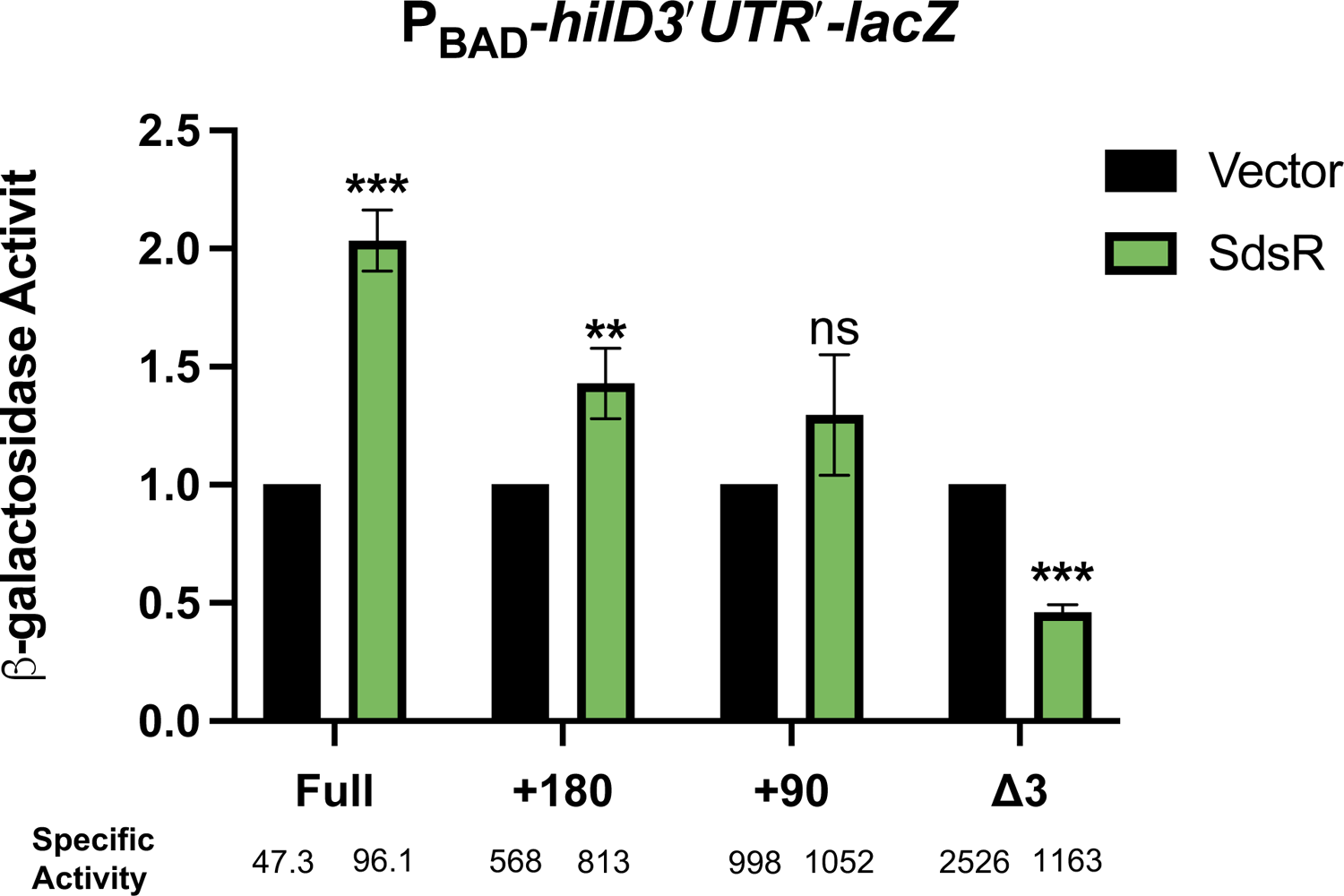
SdsR directly regulates *hilD* mRNA 3’UTR. SdsR was expressed in *E. coli* strains containing the indicated *hilD 3*′*UTR-*′*lacZ* fusions. Relative β-galactosidase units were obtained by normalizing β-galactosidase activity to vector control and reported as mean ± standard deviation. Error bars are standard deviations of the results of three independent experiments, obtained by unpaired *t* test, n=3. **, *P* < 0.005; ***, *P* < 0.0005, ns, not significant.

**Figure S3:**
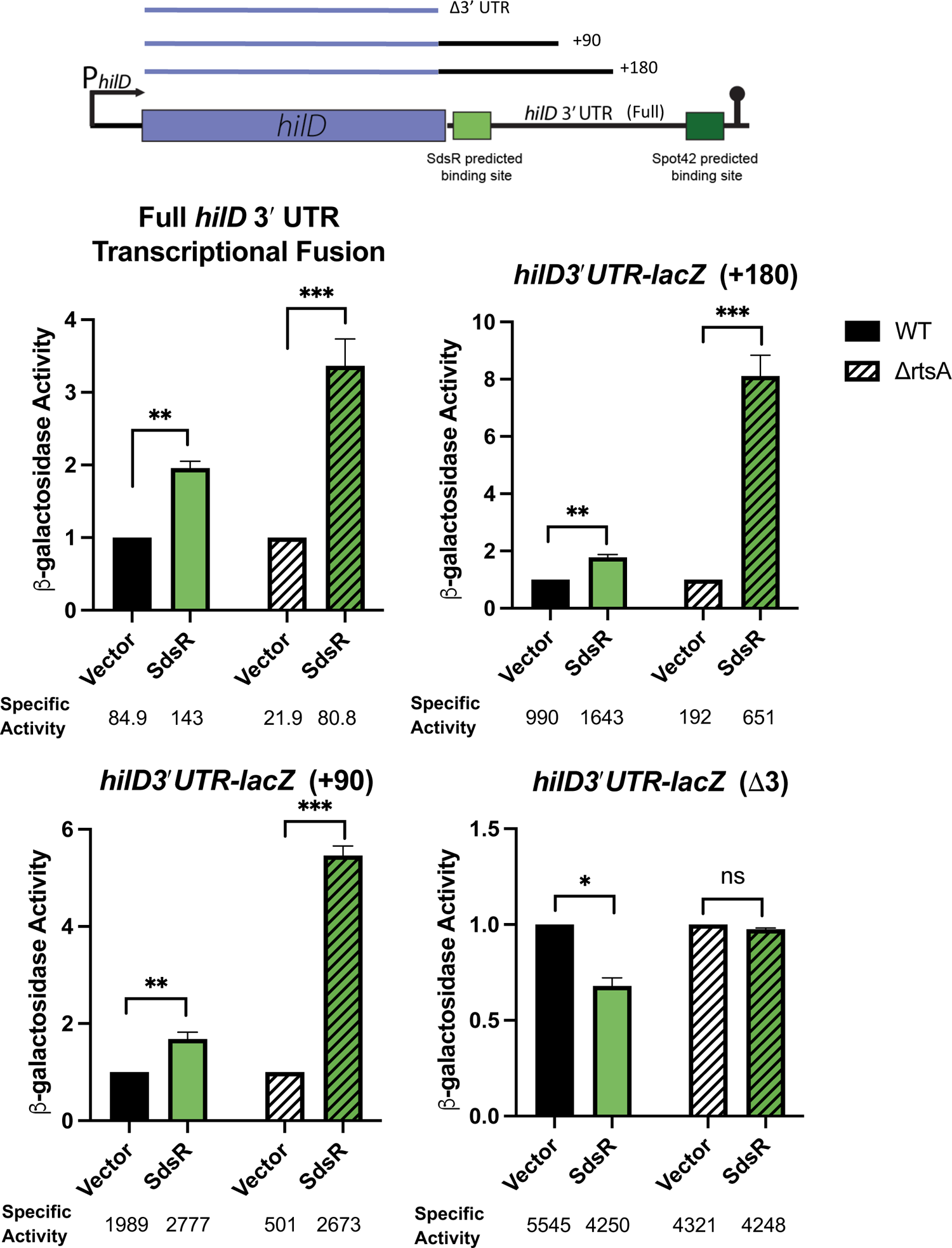
SdsR-mediated regulation of *hilD* mRNA 3′ UTR is more dramatic in *rtsA* deletion background. SdsR was expressed in strains containing the indicated *hilD 3*′*UTR-*′*lacZ* fusions in either a WT (*rtsA+*) background or a Δ*rtsA* background. Relative β-galactosidase units were obtained by normalizing β-galactosidase activity to vector control and reported as mean ± standard deviation. Error bars are standard deviations of the results of three independent experiments, obtained by unpaired *t* test, n=3. *, *P* < 0.05; **, *P* < 0.005; ***, *P* < 0.0005, ns, not significant.

**Figure S4:**
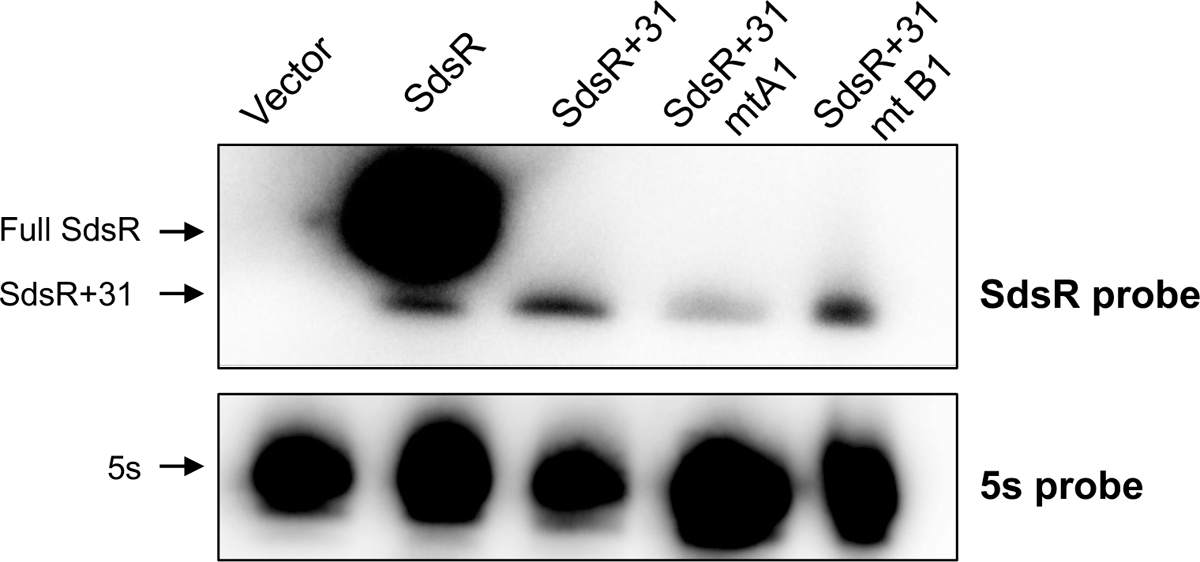
Stability of SdsR+31 variants. Stability of SdsR mutants was evaluated via northern blot using a 5′-end labelled probe corresponding to SdsR. SdsR +31 and SdsR +31 mt B1 are stable. 5S rRNA serves as a loading control.

**Figure S5.**
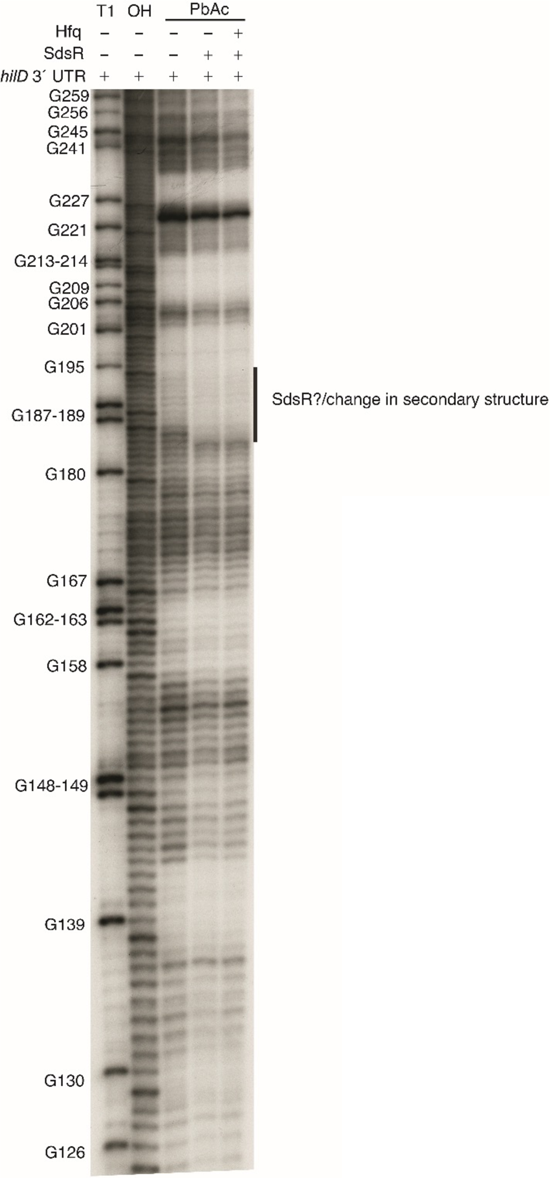
RNA footprint of a downstream fragment of *hilD* 3’UTR (+80-270): RNA footprint of a *hilD* mRNA 3′ UTR (+80-+270) alone, with SdsR, or with SdsR and Hfq.

## References

1. Acheson D, Hohmann EL. 2001. Nontyphoidal *Salmonellosis*. Clinical Infectious Diseases 32:263–269.

2. Pang T, Bhutta ZA, Finlay BB, Altwegg M. 1995. Typhoid fever and other *salmonellosis*: a continuing challenge. Trends in Microbiology 3:253–255.

3. Jones BD, Falkow S. 1996. SALMONELLOSIS: Host Immune Responses and Bacterial Virulence Determinants. Annual Review of Immunology 14:533–561.

4. Anderson CJ, Kendall MM. 2017. *Salmonella enterica* serovar Typhimurium strategies for host adaptation. Frontiers in Microbiology 8:1983–1983.

5. Galán JE. 2001. *Salmonella* Interactions with Host Cells: Type III Secretion at Work. Annual Review of Cell and Developmental Biology 17:53–86.

6. Galán JE, Collmer A. 1999. Type III secretion machines: Bacterial devices for protein delivery into host cells. Science 284:1322–1328.

7. Sturm A, Heinemann M, Arnoldini M, Benecke A, Ackermann M, Benz M, Dormann J, Hardt WD. 2011. The cost of virulence: Retarded growth of *Salmonella* Typhimurium cells expressing type III secretion system 1. PLoS Pathogens 7:e1002143–e1002143.

8. Ellermeier JR, Slauch JM. 2007. Adaptation to the host environment: regulation of the SPI1 type III secretion system in *Salmonella enterica* serovar Typhimurium. Current Opinion in Microbiology 10:24–29.

9. Golubeva YA, Sadik AY, Ellermeier JR, Slauch JM. 2012. Integrating global regulatory input into the *Salmonella* pathogenicity Island 1 type III secretion system. Genetics 190:79–90.

10. Altier C. 2005. Genetic and environmental control of *Salmonella* invasion. Journal of Microbiology 43:85–92.

11. Cott Chubiz JE, Golubeva YA, Lin D, Miller LD, Slauch JM, Chubiz JEC, Golubeva YA, Lin D, Miller LD, Slauch JM. 2010. FliZ regulates expression of the *Salmonella* pathogenicity island 1 invasion locus by controlling HilD protein activity in *Salmonella enterica* serovar Typhimurium. Journal of Bacteriology 192:6261–6270.

12. Ellermeier JR, Slauch JM. 2008. Fur regulates expression of the *Salmonella* pathogenicity island 1 type III secretion system through HilD. Journal of Bacteriology 190:476–86.

13. Lucas RL, Lee CA. 2001. Roles of *hilC* and *hilD* in regulation of *hilA* expression in *Salmonella enterica* serovar Typhimurium. Journal of Bacteriology 183:2733–2745.

14. Saini S, Ellermeier JR, Slauch JM, Rao CV. 2010. The role of coupled positive feedback in the expression of the SPI1 type three secretion system in *Salmonella*. PLoS Pathogens 6:1–16.

15. Gottesman S, McCullen CA, Guillier M, Vanderpool CK, Majdalani N, Benhammou J, Thompson KM, FitzGerald PC, Sowa NA, FitzGerald DJ. 2006. Small RNA regulators and the bacterial response to stress. Cold Spring Harb Symp Quant Biol 71:1–11.

16. Hébrard M, Kröger C, Srikumar S, Colgan A, Händler K, Hinton JCD. 2012. sRNAs and the virulence of *Salmonella enterica* serovar Typhimurium. RNA Biology 9:437–445.

17. Moll I, Afonyushkin T, Vytvytska O, Kaberdin VR, Bläsi U. 2003. Coincident Hfq binding and RNase E cleavage sites on mRNA and small regulatory RNAs. RNA 9:1308–1314.

18. De Lay N, Schu DJ, Gottesman S. 2013. Bacterial small RNA-based negative regulation: Hfq and its accomplices. The Journal of Biological Chemistry 288:7996– 8003.

19. Holmqvist E, Li L, Bischler T, Barquist L, Vogel J. 2018. Global Maps of ProQ Binding In Vivo Reveal Target Recognition via RNA Structure and Stability Control at mRNA 3′ Ends. Molecular Cell 70:971–982.e6.

20. Holmqvist E, Wright PR, Li L, Bischler T, Barquist L, Reinhardt R, Backofen R, Vogel J. 2016. Global RNA recognition patterns of post-transcriptional regulators Hfq and CsrA revealed by UV crosslinking *in vivo*. The EMBO journal 35:e201593360–e201593360.

21. Gottesman S, Storz G. 2011. Bacterial Small RNA Regulators: Versatile Roles and Rapidly Evolving Variations. Cold Spring Harbor Perspectives in Biology 3:a003798–a003798.

22. Majdalani N, Vanderpool CK, Gottesman S. 2005. Bacterial Small RNA Regulators. Critical Reviews in Biochemistry and Molecular Biology 40:93–113.

23. Papenfort K, Vanderpool CK. 2015. Target activation by regulatory RNAs in bacteria. FEMS microbiology reviews 39:362–78.

24. Morita T, Aiba H. 2011. RNase E action at a distance: Degradation of target mRNAs mediated by an Hfq-binding small RNA in bacteria. Genes and Development 25:294–298.

25. López-Garrido J, Puerta-Fernández E, Casadesús J. 2014. A eukaryotic-like 3′ untranslated region in *Salmonella enterica hilD* mRNA. Nucleic Acids Research 42:5894–5906.

26. El Mouali Y, Gaviria-Cantin T, Sánchez-Romero MA, Gibert M, Westermann AJ, Vogel J, Balsalobre C. 2018. CRP-cAMP mediates silencing of *Salmonella* virulence at the post-transcriptional level. PLoS Genetics 14.

27. Kim K, Palmer AD, Vanderpool CK, Slauch JM. 2019. The small RNA PinT contributes to *phoP*-mediated regulation of the *Salmonella* Pathogenicity Island 1 type III secretion system in *Salmonella enterica* serovar Typhimurium. Journal of Bacteriology 201.

28. Kim K, Golubeva YA, Vanderpool CK, Slauch JM. 2019. Oxygen-Dependent Regulation of SPI1 Type Three Secretion System by Small RNAs in *Salmonella enterica* serovar Typhimurium. Molecular Microbiology mmi.14174-mmi.14174.

29. Cakar F, Golubeva YA, Vanderpool CK, Slauch JM. The sRNA MicC downregulates *hilD* translation to control the SPI1 T3SS in *Salmonella enterica* serovar Typhimurium. Journal of Bacteriology 0:JB.00378-21.

30. Padalon-Brauch G, Hershberg R, Elgrably-Weiss M, Baruch K, Rosenshine I, Margalit H, Altuvia S. 2008. Small RNAs encoded within genetic islands of *Salmonella* typhimurium show host-induced expression and role in virulence. Nucleic Acids Research https://doi.org/10.1093/nar/gkn050.

31. Sharma CM, Vogel J. 2009. Experimental approaches for the discovery and characterization of regulatory small RNA. Current Opinion in Microbiology 12:536– 546.

32. Gaviria-Cantin T, Mouali YE, Guyon SL, Römling U, Balsalobre C. 2017. Gre factors-mediated control of *hilD* transcription is essential for the invasion of epithelial cells by *Salmonella enterica* serovar Typhimurium. PLOS Pathogens 13:e1006312.

33. Lee CA, Jones BD, Falkow S. 1992. Identification of a *Salmonella* Typhimurium invasion locus by selection for hyperinvasive mutants. Proc Natl Acad Sci USA 89:1847.

34. Song J-WW, Woo J-MM, Jung GY, Bornscheuer UT, Park J-BB. 2016. 3′-UTR engineering to improve soluble expression and fine-tuning of activity of cascade enzymes in *Escherichia coli*. Scientific Reports 6:29406–29406.

35. Mandin P, Gottesman S. 2009. A genetic approach for finding small RNAs regulators of genes of interest identifies RybC as regulating the DpiA/DpiB two-component system. Molecular Microbiology 72:551–565.

36. Lopez PJ, Marchand I, Joyce SA, Dreyfus M. 1999. The C-terminal half of RNase E, which organizes the *Escherichia coli* degradosome, participates in mRNA degradation but not rRNA processing in vivo. Molecular Microbiology 33:188–199.

37. Pfeiffer V, Sittka A, Tomer R, Tedin K, Brinkmann V, Vogel J. 2007. A small non-coding RNA of the invasion gene island (SPI-1) represses outer membrane protein synthesis from the *Salmonella* core genome. Molecular Microbiology 66:1174– 1191.

38. Fröhlich KS, Fröhlich F, Haneke K, Papenfort K, Org Vogel J”. 2016. The target spectrum of SdsR small RNA in *Salmonella*. Nucleic Acids Research 44:10406– 10422.

39. Polayes DA, Rice PW, Garner MM, Dahlberg JE. 1988. Cyclic AMP-cyclic AMP receptor protein as a repressor of transcription of the *spf* gene of *Escherichia coli*. Journal of Bacteriology 170:3110–4.

40. Azam MS, Vanderpool CK. 2020. Translation inhibition from a distance: the small RNA SgrS silences a ribosomal protein S1-dependent enhancer. Mol Microbiol 114:391–408.

41. Bianco CM, Fröhlich KS, Vanderpool CK. 2019. Bacterial Cyclopropane Fatty Acid Synthase mRNA Is Targeted by Activating and Repressing Small RNAs. Journal of Bacteriology 201:e00461–19.

42. Bobrovskyy M, Vanderpool CK. 2016. Diverse mechanisms of post-transcriptional repression by the small RNA regulator of glucose-phosphate stress. Molecular Microbiology 99:254–273.

43. Azam MS, Vanderpool CK. 2018. Translational regulation by bacterial small RNAs via an unusual Hfq-dependent mechanism. Nucleic Acids Research 46:2585–2599.

44. Link TM, Valentin-Hansen P, Brennan RG. 2009. Structure of *Escherichia coli* Hfq bound to polyriboadenylate RNA. Proceedings of the National Academy of Sciences 106:19292–19297.

45. Vazquez-Torres A, Jones-Carson J, Bäumler AJ, Falkow S, Valdivia R, Brown W, Le M, Berggren R, Parks WT, Fang FC. 1999. Extraintestinal dissemination of Salmonella by CD18-expressing phagocytes. 6755. Nature 401:804–808.

46. Anderson CJ, Kendall MM. 2017. *Salmonella enterica*, serovar Typhimurium strategies for host adaptation. Frontiers in Microbiology 8:1983–1983.

47. Ruiz de los Mozos I, Vergara-Irigaray M, Segura V, Villanueva M, Bitarte N, Saramago M, Domingues S, Arraiano CM, Fechter P, Romby P, Valle J, Solano C, Lasa I, Toledo-Arana A. 2013. Base Pairing Interaction between 5′- and 3′-UTRs Controls *icaR* mRNA Translation in *Staphylococcus aureus*. PLoS Genetics 9:e1004001–e1004001.

48. Bronesky D, Desgranges E, Corvaglia A, François P, Caballero CJ, Prado L, Toledo-Arana A, Lasa I, Moreau K, Vandenesch F, Marzi S, Romby P, Caldelari I. 2019. A multifaceted small RNA modulates gene expression upon glucose limitation in *Staphylococcus aureu*s. EMBO J 38:e99363.

49. Narm K-E, Kalafatis M, Slauch JM. 2020. HilD, HilC, and RtsA Form Homodimers and Heterodimers To Regulate Expression of the *Salmonella* Pathogenicity Island I Type III Secretion System. Journal of Bacteriology 202.

50. Krol J, Loedige I, Filipowicz W. 2010. The widespread regulation of microRNA biogenesis, function and decay. Nat Rev Genet 11:597–610.

51. O’Brien J, Hayder H, Zayed Y, Peng C. 2018. Overview of MicroRNA Biogenesis, Mechanisms of Actions, and Circulation. Frontiers in Endocrinology 9.

52. Datsenko KA, Wanner BL. 2000. One-step inactivation of chromosomal genes in *Escherichia coli* K-12 using PCR products. Proceedings of the National Academy of Sciences of the United States of America 97:6640–5.

53. Maloy SR, Stewart VJ, Taylor RK. 1996. Genetic analysis of pathogenic bacteria: a laboratory manual. Cold Spring Harbor Laboratory Press, Plainview, N.Y.

54. Guillier M, Gottesman S. 2006. Remodeling of the *Escherichia coli* outer membrane by two small regulatory RNAs. Molecular Microbiology 59:231–247.

55. Busch A, Richter AS, Backofen R. 2008. IntaRNA: efficient prediction of bacterial sRNA targets incorporating target site accessibility and seed regions. Bioinformatics 24:2849–2856.

56. Slauch JM, Silhavy TJ. 1991. cis-Acting *ompF* mutations that result in OmpR-dependent constitutive expression. Journal of Bacteriology 173:4039–4048.

57. Desnoyers G, Morissette A, Prévost K, Massé E. 2009. Small RNA-induced differential degradation of the polycistronic mRNA *iscRSUA*. EMBO Journal 28:1551–1561.

58. Aiba H, Adhya S, de Crombrugghe B. 1981. Evidence for two functional gal promoters in intact *Escherichia coli* cells. Journal of Biological Chemistry 256:11905–11910.

59. Fröhlich KS, Papenfort K, Fekete A, Vogel J. 2013. A small RNA activates CFA synthase by isoform-specific mRNA stabilization. The EMBO Journal 32:2963– 2979.

60. Lin D, Rao CV, Slauch JM. 2008. The *Salmonella* SPI-1 type three secretion system responds to periplasmic disulfide bond status via the flagellar apparatus and the RcsCDB system. Journal of Bacteriology 190:87–97.

61. Ellermeier CD, Slauch JM. 2003. RtsA and RtsB coordinately regulate expression of the invasion and flagellar genes in *Salmonella enterica* serovar Typhimurium. Journal of Bacteriology 185:5096–108.

62. Ellermeier CD, Ellermeier JR, Slauch JM. 2005. HilD, HilC and RtsA constitute a feed forward loop that controls expression of the SPI-1 type three secretion system regulator hilA in Salmonella enterica serovar Typhimurium. Molecular Microbiology 57:691–705.

63. Cott Chubiz JE, Golubeva YA, Lin D, Miller LD, Slauch JM, Chubiz JEC, Golubeva YA, Lin D, Miller LD, Slauch JM. 2010. FliZ regulates expression of the Salmonella Pathogenicity Island 1 invasion locus by controlling HilD protein activity in *Salmonella enterica* serovar Typhimurium. Journal of Bacteriology 192:6261–6270.

64. Stanley TL, Ellermeier CD, Slauch JM. 2000. Tissue-specific gene expression identifies a gene in the lysogenic phage Gifsy-1 that affects *Salmonella enterica* serovar Typhimurium survival in Peyer’s patches. Journal of Bacteriology 182:4406–4413.

